# CHESS 3: an improved, comprehensive catalog of human genes and transcripts based on large-scale expression data, phylogenetic analysis, and protein structure

**DOI:** 10.1101/2022.12.21.521274

**Authors:** Ales Varabyou, Markus J. Sommer, Beril Erdogdu, Ida Shinder, Ilia Minkin, Kuan-Hao Chao, Sukhwan Park, Jakob Heinz, Christopher Pockrandt, Alaina Shumate, Natalia Rincon, Daniela Puiu, Martin Steinegger, Steven L. Salzberg, Mihaela Pertea

## Abstract

The original CHESS database of human genes was assembled from nearly 10,000 RNA sequencing experiments in 53 human body sites produced by the Genotype-Tissue Expression (GTEx) project, and then augmented with genes from other databases to yield a comprehensive collection of protein-coding and noncoding transcripts. The construction of the new CHESS 3 database employed improved transcript assembly algorithms, a new machine learning classifier, and protein structure predictions to identify genes and transcripts likely to be functional and to eliminate those that appeared more likely to represent noise. The new catalog contains 41,356 genes on the GRCh38 reference human genome, of which 19,839 are protein-coding, and a total of 158,377 transcripts. These include 14,863 novel protein-coding transcripts. The total number of transcripts is substantially smaller than earlier versions due to improved transcriptome assembly methods and to a stricter protocol for filtering out noisy transcripts. Notably, CHESS 3 contains all of the transcripts in the MANE database, and at least one transcript corresponding to the vast majority of protein-coding genes in the RefSeq and GENCODE databases. CHESS 3 has also been mapped onto the complete CHM13 human genome, which gives a more-complete gene count of 43,773 genes and 19,968 protein-coding genes. The CHESS database is available at http://ccb.jhu.edu/chess.

## Background

With the first release in 2021 of a truly complete human genome, designated CHM13 [1], the scientific community now has the opportunity to complete the Human Genome Project by identifying not only the sequence, but also all of the genes in the genome. The T2T

Consortium’s assembly reported 2,226 additional copies of known human genes and a total of 63,494 genes, including 19,969 protein-coding loci with 86,245 transcripts. That annotation was produced by mapping the annotation from GENCODE v35 [2] onto the CHM13 assembly, followed by using Liftoff [3] to identify extra gene copies. Thus although the CHM13 gene list is more complete than the corresponding GRCh38 annotation, it does not include all of the genes in RefSeq [4], CHESS [5], FANTOM [6], APPRIS [7], or other human gene databases.

The CHESS human gene catalog, first published in 2018 [5], is an effort to provide a comprehensive database of human genes that includes all protein-coding and noncoding genes. Unlike other efforts, the basis of nearly all CHESS genes is direct experimental evidence from RNA sequencing experiments, in particular the large-scale Genotype-Tissue Expression (GTEx) project, which has generated thousands of deep RNA sequencing datasets from hundreds of individuals and dozens of tissue types [8]. The construction of CHESS begins with a large-scale assembly of all of these experiments, producing millions of transcripts that are then filtered to generate the final database. As described below, this process means that almost every gene in CHESS can be linked directly to experimental evidence for that gene’s expression. To ensure its completeness, and because GTEx does not capture 100% of human genes, we identify and add to CHESS any well-supported genes in other databases that were not assembled from the GTEx data.

Despite decades of effort, the primary human gene databases still do not agree on the precise number or structure of human genes, reflecting the difficulty of this task. The latest release of CHESS includes substantially improved transcriptome assembly methods, a novel machine learning strategy to identify reliable introns, and new validation steps based on protein structure prediction, but nonetheless it is not expected to be the final, authoritative list of human genes. In an effort to make CHESS as complete as possible, we augmented the assembled gene list by ensuring that it contains all of the genes in the MANE database, a recently-developed (but still incomplete) catalog that has one high-quality transcript for nearly all protein-coding genes, and for which RefSeq and GENCODE agree precisely on the transcript boundaries and on the coding sequence [9].

CHESS 3 takes a stricter approach to including genes and transcripts than other human gene catalogs, including previous versions of CHESS. In particular, we do not include in the primary database any gene or transcript that appears to be non-functional, although we do provide separate sets of assembled transcripts for users who want them. This strategy means that aberrant transcripts, such as those created by erroneous splicing or those that create truncated and non-functional proteins, are not included in CHESS. Other catalogs include thousands of these transcripts, sometimes tagged to indicate they are non-functional, but sometimes merely included without any such warning. A growing body of evidence suggests that many alternative splicing events do not produce functional proteins [10]. We have described how these non-functional transcripts, which usually occur at very low expression levels, are likely to confuse analysis software and produce misleading results [11], and annotation databases will be improved by excluding them.

## Results

The CHESS 3 catalog is based principally on direct evidence from RNA-sequencing experiments, in particular the GTEx collection of transcripts from 53 body sites and hundreds of individuals [8]. All transcripts were processed through a complex alignment, assembly, and filtering process (see Methods), which eliminated millions of transcript fragments representing noise.

CHESS 3 contains 19,839 protein-coding genes with a total of 99,202 transcripts, approximately 5 transcripts per gene. If we exclude duplicate amino-acid sequences, the number of distinct protein sequences produced from these transcripts is 73,767 (Table **1**). In total, including noncoding transcripts, CHESS 3 has 158,377 transcripts on the primary chromosomes. (Note that GRCh38 also has several hundred alternative scaffolds containing thousands of annotated genes, the vast majority of which are duplicates. For consistency, we are only counting genes placed on the primary chromosomes in this discussion.)

**Table 1.** Total number of genes and protein-coding isoforms in current versions of CHESS, RefSeq, and GENCODE. Genes are counted on the primary chromosomes and unplaced scaffolds from the human reference genome GRCh38, excluding the alternative scaffolds. Pseudogenes, VDJ segments, and C regions are not included in the totals shown in the final column.

| **Table 1.** Total number of genes and protein-coding isoforms in current versions of CHES3, RefSeq, and GENCODE. Genes are counted on the primary chromosomes and unplaced scaffolds from the human reference genome GRCh38, excluding the alternative scaffolds. Pseudogenes, VDJ segments, and C regions are not included in the totals shown in the final column. |  |  |  |  |
| --- | --- | --- | --- | --- |
| Database | Number of protein-coding gene loci | Number of protein-coding transcripts | Number of distinct protein sequences | Number of gene loci (all types) |
| CHES3 v3 | 19,839 | 99,202 | 73,767 | 41,356 |
| RefSeq v110 | 19,884 | 129,740 | 88,662 | 43,380 |
| GENCODE v41 | 19,419 | 110,309 | 92,968 | 46,181 |

By comparison, the latest version of GENCODE (release 41) contains 19,419 protein-coding gene loci on the primary chromosomes, containing 110,309 protein-coding transcripts that encode 92,968 distinct protein sequences. RefSeq (release 110) has 19,884 protein-coding genes and 129,740 protein-coding transcripts, encoding 88,662 different protein sequences (Table **1**).

In CHESS 3, all transcripts at protein-coding loci are required to have valid open reading frames (ORFs) corresponding to the protein sequences encoded by those transcripts. These are represented as CDS features in the annotation file. Any alternative splice variant or isoform that does not produce a functional protein is considered to be transcriptional noise, and is not annotated as a transcript. RefSeq follows a similar strategy, where nearly every transcript (with a few exceptions) at a protein-coding locus contains a valid ORF. In contrast, GENCODE contains thousands of transcripts at protein-coding loci that do not encode functional proteins for a variety of reasons, which are indicated by tags such as “retained intron” (33,750 transcripts) or “nonsense mediated decay” (20,933 transcripts).

In addition to removing assembled transcripts that did not contain a valid ORF, as part of the CHESS 3 refinement process we evaluated the relative lengths of all protein sequences at each locus. We assume that severely truncated proteins are highly unlikely to be functional, and therefore the transcripts encoding them should, with few exceptions, be classified as noise and removed. Based on analysis of protein lengths in RefSeq, we chose a threshold of one-fifth the maximum length at a locus, and any protein shorter than that was considered non-functional and removed from CHESS, unless there was independent evidence that it was functional (see Methods).

Note that the CHESS 3 data release includes a separate catalog of transcripts that were assembled from the GTEx collection, but that were filtered out because they lack a valid translation or because the translated protein is too short. This provides a resource for those who wish to explore transcriptional noise itself, or to mine the data looking for transcripts that might be re-classified as functional.

To illustrate the variability in protein lengths in different annotation databases, consider the Titin (TTN) protein, the longest in the human genome at 35,991 amino acids (aa). GENCODE v41 includes 15 protein-coding transcripts for Titin, ranging from 48 to 35,991aa, with eight isoforms shorter than 1000aa (Table **2**). The transcripts shorter than 1000aa at this locus are almost certainly non-functional, and indeed GENCODE annotates them as having incomplete coding sequences at either the 5’ end, the 3’ end, or both.

**Table 2.** Isoforms of the protein-coding gene Titin (TTN), gene ID ENSG00000155657.29, in GENCODE v41, showing the length of the annotated proteins for each of 15 isoforms. Isoforms whose lengths are marked with * are also present in both CHESS and RefSeq.

| Transcript ID | Translated protein length (aa) |
| --- | --- |
| ENST00000412264.1 | 48 |
| ENST00000448510.2 | 172 |
| ENST00000436599.1 | 213 |
| ENST00000425332.2 | 240 |
| ENST00000426232.5 | 255 |
| ENST00000634225.1 | 353 |
| ENST00000446966.1 | 372 |
| ENST00000414766.5 | 962 |
| ENST00000360870.10 | 5604* |
| ENST00000460472.6 | 26926* |
| ENST00000359218.10 | 27051* |
| ENST00000342175.11 | 27118* |
| ENST00000342992.11 | 33423* |
| ENST00000591111.5 | 34350* |
| ENST00000589042.5 | 35991* |

By contrast, RefSeq’s (v110) 22 isoforms of the Titin gene range in length from 23564 to 35991 aa, with one shorter isoform at 5604aa. That relatively short isoform, present in GENCODE as well, has been subject to experiments that show that it is both transcribed and translated, and that also demonstrate its possible function [12]. CHESS has 8 isoforms with the shortest also at 5604aa. Worth noting here is that no isoform shorter than 1000aa exists in either RefSeq or CHESS, while all of the longer isoforms in GENCODE, including the 5604aa variant, are in both RefSeq and CHESS.

To consider just one more example, in RefSeq the protein with the greatest ratio between longest and shortest isoforms is AHNAK, a 5890aa protein that has a 149aa isoform. The unusually short isoform has been shown experimentally to fulfill a self-regulatory role in muscle [13], thus despite the very short length, there is independent evidence to support it. While RefSeq and CHESS contain only this one short isoform of AHNAK, GENCODE contains six others, with lengths ranging from 85 to 149aa, in addition to the long isoform at 5890aa. Only the 149aa and 5890aa isoforms are supported by experimental evidence.

Extreme variation in length is seen among many other annotated transcripts in GENCODE, where we found 4089 protein-coding genes that have an isoform whose length is <10% of the length of the longest isoform, and 7269 protein-coding genes that have an isoform whose length is <20% of the longest isoform. In contrast, both RefSeq and CHESS contain far fewer protein-coding genes for which the isoforms vary so dramatically in length. RefSeq contains just 79 genes for which the longest isoform is at least 10 times the length of the shortest, and 333 genes where the longest isoform is at least 5 times longer than the shortest. CHESS only has 4 such genes: the Titin and AHNAK genes mentioned above, and two genes (IQSEC2 and SYNE1) from the MANE database that are tagged as special isoforms of clinical significance.

Also worth noting is that the shortest protein sequence (RPL41, ribosomal protein L41) in RefSeq is 25aa long, while GENCODE contains 1259 protein isoforms that are shorter than 25aa, including 20 annotated CDS features whose length is just 1aa. CHESS contains only 14 protein isoforms shorter than 25aa.

### Inclusion of MANE transcripts in CHESS

The creators of RefSeq and GENCODE have released a high-quality collection of protein-coding transcripts called MANE (Matched Annotation between NCBI and EMBL-EBI), which they have described as a “universal standard” for human gene annotation [9]. MANE is an effort to annotate one transcript for each human protein-coding gene for which RefSeq and GENCODE agree perfectly, including the 5’ and 3’ boundaries of transcription, all exon and intron boundaries, as well as the coding sequence. The current release of MANE (v1.0) has 19,062 proteins and 19,120 transcripts, with the extra 58 transcripts included because of their clinical significance. MANE does not include any noncoding genes.

Because MANE is both high-quality and stable, we wanted to ensure that every transcript in MANE was also included in CHESS 3. After comparing our near-final set of transcripts to MANE, we found that nearly all of them had a near-perfect match to one CHESS transcript, although a small number had differences in the precise boundaries at the beginning and end of transcription. We then edited the 5’ and 3’ boundaries so that one CHESS transcript matches MANE perfectly for all 19,120 of the MANE transcripts, with no exceptions.

#### Novel protein-coding genes in CHESS 2 and CHESS 3

We reported previously [5] that the CHESS database (v2.2) had 224 novel proteins that were missing entirely from both RefSeq and GENCODE. We investigated the current releases of both databases (v110 and v41 respectively) and found 5 of the previously novel protein-coding genes that are now included in GENCODE, RefSeq, and MANE, shown in Table 3.

**Table 3.** Protein-coding genes that were novel in CHESS release 1 and 2, and that are now part of the GENCODE, RefSeq, and MANE databases.

| **Table 3.** Protein-coding genes that were novel in CHES release 1 and 2, and that are now part of the GENCODE, RefSeq, and MANE databases. |  |  |  |  |
| --- | --- | --- | --- | --- |
| CHES 3 ID | Coordinates (chr:position) | Gene name | GENCODE ID | RefSeq ID |
| CHS.3390 | 1:158125775-158130906 | SMIM42 | ENSG00000288460.1 | NM_001395415.1 |
| CHS.7402 | 10:122657410-122679509 |  | ENSG00000286135.1 | NM_001364461.3 |
| CHS.8823 | 11:59880266-59896469 | OOSP3 | ENSG00000285231.2 | NM_001395255.1 |
| CHS.11637 | 12:51813540-51814957 | TMDD1 | ENSG00000284730.2 | NM_001386737.1 |
| CHS.58949 | X:149414886-149415495 |  | ENSG00000287585.2 | NM_001395872.1 |

Every protein-coding gene locus in CHESS 3 either matches or overlaps at least one transcript in either RefSeq or GENCODE, however there are many protein-coding transcripts that are unique to each of the databases. We considered a pair of transcripts a match if all introns matched precisely; using this criterion, 14,863 out of 99,201 protein-coding transcripts in CHESS 3 are unique to CHESS (**Figure 1**). Another 46,585 of those transcripts are shared by all 3 databases, while 32,882 are shared by CHESS and RefSeq only, and 4871 are shared by CHESS and GENCODE only. RefSeq and GENCODE share 658 protein-coding transcripts that are not in CHESS.

**Figure 1.**
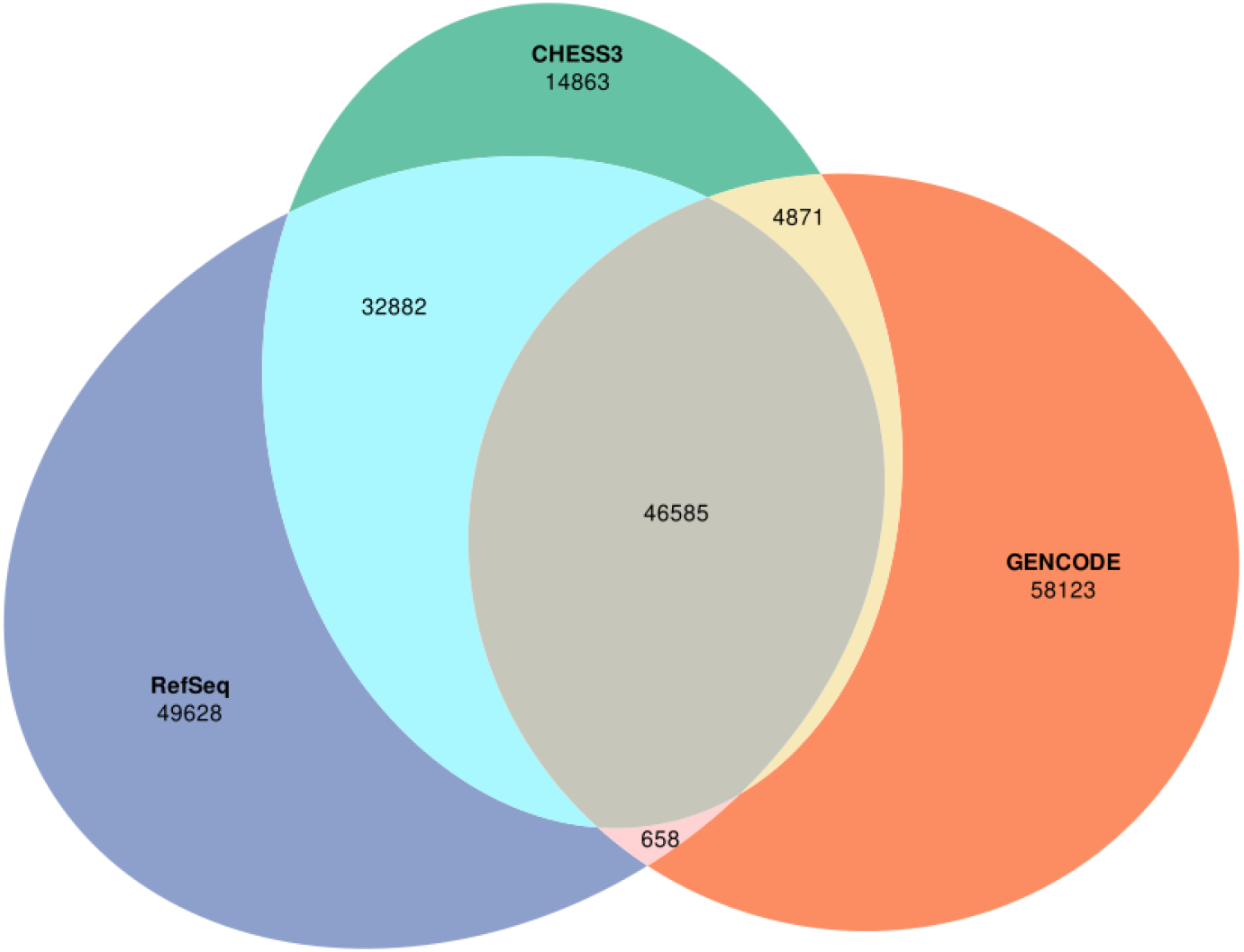
Overlap between the protein-coding transcripts in CHESS 3, RefSeq v110, and GENCODE v41. Transcripts were considered matching if all of their introns match precisely.

#### Comparisons to RefSeq and GENCODE

In earlier releases of CHESS, we made a conscious decision to include all protein-coding gene loci (although not all transcripts) from RefSeq and GENCODE in the CHESS database. However, upon closer scrutiny, we discovered that some of these genes are likely not true protein-coding genes, but instead are legacy annotations from earlier versions of those databases. Both RefSeq and GENCODE have removed many of their genes over the years, but a few genes with very weak evidence still remain.

We compared CHESS 3 to all RefSeq (v110) protein-coding genes, and identified 46 loci that are missing from CHESS. All of these have names beginning with “LOC,” indicating that their function is unknown, and each is annotated with an “XM” designation by RefSeq, which means it is a low-confidence annotation (as opposed to “NM” genes, which are considered high-confidence). Many are also contained within the introns of other genes; for example, LOC107984876 (XM_047434996.1) is contained within exon 4 of the protein-coding gene LMF1 on chromosome 16, and a search of its putative sequence has no hits outside primates. This evidence, combined with the fact that we did not assemble these genes from the GTEx data, led us to decide not to add them to CHESS 3.

Overall, CHESS, RefSeq, and GENCODE are in closer agreement today than they were in 2018, when the previous major release (2.0) of CHESS appeared. **Figure 1** illustrates the overlap between protein-coding transcripts among all three databases. Compared to the 2018 versions of CHESS (v2.2), GENCODE (v28), and RefSeq (v108), the number of transcripts shared among all three databases has increased substantially, from 36,943 to 46,585. Although still very high, the number of transcripts unique to any of the three databases has declined from 189,184 to 122,614, largely due to the decline in the number of protein-coding transcripts in CHESS.

#### Protein structure predictions for CHESS 3

We used the AlphaFold2 [14] and ColabFold [15] programs to predict the three-dimensional structure of all but the largest protein isoforms in CHESS 3, making it the only human annotation database currently to include structure predictions for most of its proteins. Specifically, we used ColabFold (version d6b06) to predict the structures for >230,000 transcripts from a preliminary version of CHESS 3, which was a superset of the final database. These included all proteins in CHESS 3 shorter than 1000aa. We then collected predictions for longer proteins from the AlphaFold Protein Structure Database v3 [16] that exactly matched isoforms in CHESS 3. This added 3302 structures, including predictions for selected isoforms as long as 2700aa. The <u>isoform.io</u> v1.2 database contains structures for 91,589 CHESS 3 transcripts representing 70,158 unique isoforms at 19,569 protein-coding loci in CHESS 3. In total, structures are predicted for >95% of all CHESS 3 proteins covering >98% of all human protein-coding loci. All protein structures are freely available for searching or download at isoform.io, which contains cross-references to CHESS, RefSeq, GENCODE, and MANE for each structure.

We evaluated the 14,683 protein-coding transcripts that are unique to CHESS 3 to identify those that have unique protein sequences and are highly expressed as well. We restricted our search to multi-exon protein-coding genes that had a protein-coding sequence that was non-identical to any other annotated protein. We also required that these novel transcripts had a cumulative TPM of >1000 across all GTEx samples. Most important, we searched for transcripts where the novel proteins accounted for >50% of the total expression across all samples. These criteria yielded 261 genes with novel protein isoforms, two of which are shown in Figure 2.

**Figure 2.**
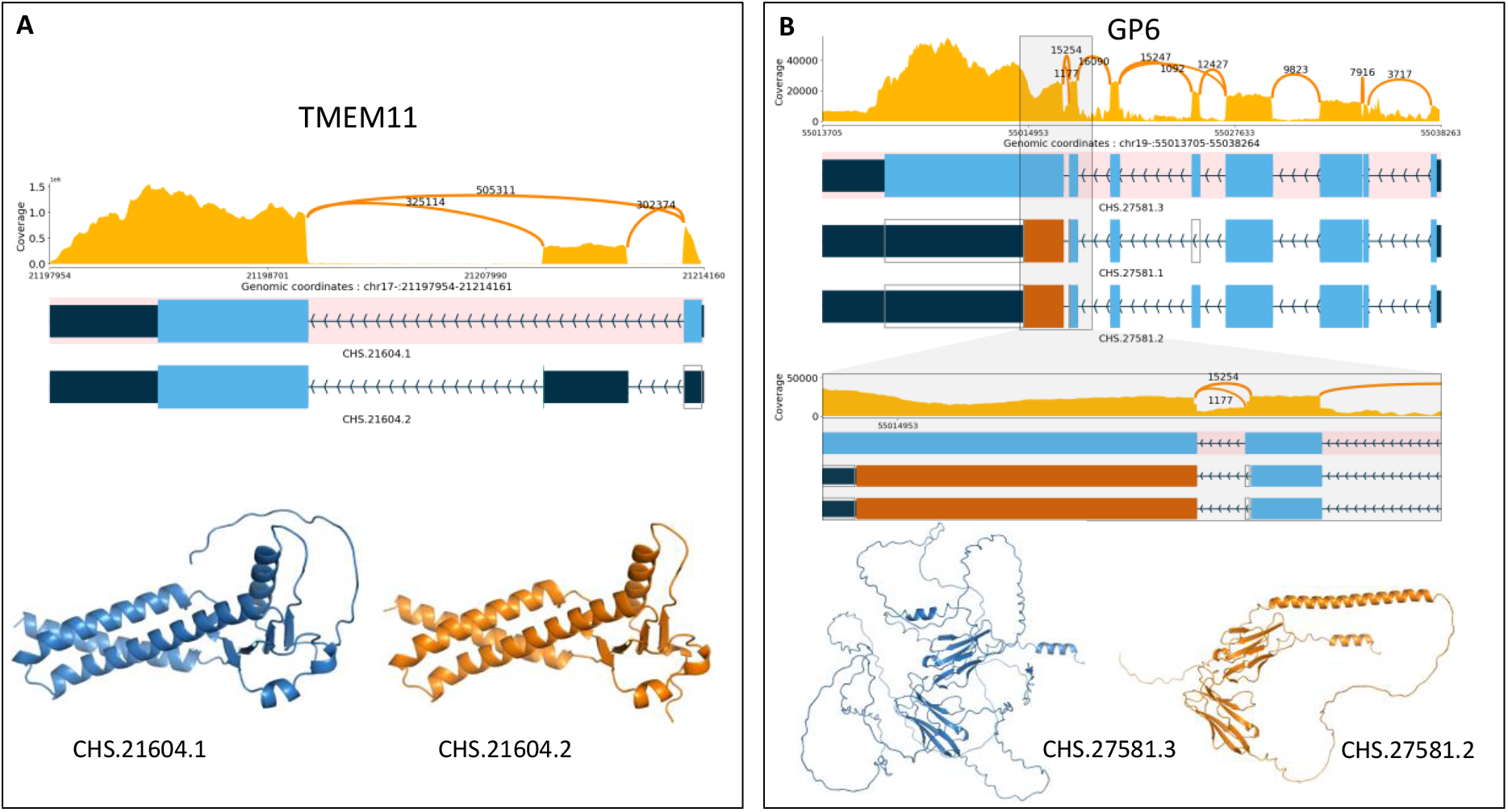
Expression levels, exon-intron structures, and protein structures for (A) TMEM11 and (B) GP6. The upper panel is a ‘Sashimi’ plot showing the total depth of RNA-seq read coverage across all tissues, with labeled arcs showing the number of spliced reads supporting each possible intron. Below that are the exon-intron structures, with the MANE isoform at the top, highlighted in pink. Protein-coding regions of exons are shown in blue and orange, where orange indicates sequence that is in a different reading frame from the MANE isoform. **(A)** The alternative isoform for TMEM11, CHS.21604.2, is unique to CHESS. The bottom shows the protein structure of both isoforms as predicted by ColabFold/Alphafold2, showing that the extra sequence in the MANE isoform is an unstructured loop. **(B)** Similar plots for 3 isoforms of GP6. An additional panel shows a zoomed-in view of the region spanning the last intron, where 15,254 spliced reads support the longer intron (in CHS.27581.1 and CHS.27581.2), while 1177 reads support the shorter intron used in the MANE isoform, CHS.27581.3. The structures at bottom show that the MANE isoform (left) has a highly disordered structure, which explains its low pLDDT score of 49.3, while the CHESS isoform on the right, which is also the highest-expressed transcript at this locus, has a score of 74.5.

The novel TMEM11 isoform shown in Figure 2A is slightly shorter than the canonical (MANE) protein, caused by an additional exon that shifts the start codon downstream. ColabFold assigns a pLDDT score of 78.6 to novel CHESS protein, versus the substantially lower score of 68.3 for the longer MANE protein, whose lower score is due to the presence of an unstructured loop. This suggests that the novel isoform might function more effectively, but answering this question will require targeted experiments. Figure 2B shows the exon-intron structures for three isoforms of GP6, where the MANE isoform has a much lower-scoring structure than the other two CHESS isoforms. The only difference between the MANE transcript and CHS.27581.2 is a 4-base shift in the last intron. The shorter intron (MANE) yields a protein that is 281aa longer (620aa versus 339aa), but the AlphaFold2 result indicates that the additional sequence is entirely unstructured, resulting in a dramatically lower pLDDT score of 49.3, versus 74.5 for the longer CHS.27581.2 protein. In addition, the longer intron (CHS.27581.2) has 13 times deeper support in spliced reads, as shown in the Sashimi plots. Note that both RefSeq and GENCODE contain isoforms matching CHS.27581.1 and CHS.27581.2. The full list of 261 novel, highly-expressed protein-coding transcripts, along with sashimi plots similar to Figure 2, is available in Supplementary **Table S1**.

#### Noncoding genes and transcripts

CHESS 3 contains 17,623 lncRNAs which encompass 34,708 transcripts, as well as many other types of noncoding transcripts (**Table 4**). RefSeq has 17,793 lncRNAs containing 29,048 transcripts, while GENCODE has 19,095 lncRNA loci and 53,216 transcripts, many more than either CHESS or RefSeq.

**Table 4.** The number of genes and transcripts on the primary chromosomes (excluding alternative scaffolds) of GRCh38 in the CHESS 3, RefSeq v110, and GENCODE v41 catalogs. lncRNA: long noncoding RNA gene; snoRNA: small nucleolar RNA; snRNA: small nuclear RNA; rRNA: ribosomal RNA; tRNA: transfer RNA.

|  |  |  |  |
| --- | --- | --- | --- |
| Gene type | CHESS | RefSeq | GENCODE |
| Protein-coding genes | 19,839 | 19,884 | 19,419 |
| lncRNA genes | 17,623 | 17,793 | 18,041 |
| microRNA | 1,914 | 1,914 | 1,879 |
| Transcript type |  |  |  |
| Protein-coding transcript | 99,202 | 129,740 | 169,195* |
| lncRNA transcript | 34,708 | 29,048 | 53,216 |
| pseudogene | 16,572 | 15,357 | 20,234 |
| tRNA | 453 | 453 | 535 |
| snoRNA | 1,195 | 1,195 | 942 |
| snRNA | 153 | 153 | 1,901 |
| rRNA | 40 | 40 | 47 |
| VDJ segments | 565 | 565 | 200 |
| Other | 213 | 12,493 | 3,822 |
| Total transcripts | **158,377** | **191,917** | **251,236** |
| *Protein-coding transcripts for GENCODE include all transcripts for genes with the biotype protein_coding, including those with tags such as "retained_intron," "mRNA_start_NF," "nonsense_mediated_decay," and others. |  |  |  |

The number of reported RNA genes has grown dramatically in recent years, with catalogs such as NONCODE [17], LNCipedia [18], lncRNAKB [19], and RNAcentral [20] containing a wide variety of gene counts. For example, as of mid-2021 NONCODE V6 had 173,112, LNCipedia had 127,802, and lncRNAKB had 77,199 human lncRNAs. Most of the lncRNAs currently annotated in these various databases represent computational predictions, and it is not known how many of them are truly genes rather than transcriptional noise. As we reported in the original description of CHESS [21], ∼98% of the transcripts initially assembled from the GTEx data appeared to be noise, and the vast majority of these were present at very low expression levels. Others have recently argued that most lncRNAs are likely to be nonfunctional, for multiple scientific reasons [22]. For CHESS 3, we attempted to use stricter criteria for including a lncRNA than for a protein, but the filtering task is made much more difficult by the fact that lncRNAs do not have open reading frames, making it harder to find sequence conservation in other species that would increase our confidence that the lncRNA is functional.

### Conservation of introns across species

To evaluate the consistency of annotation in light of evolutionary conversation, we used a 30-species alignment [23] that contained 27 primates (including human) plus mouse, dog, and armadillo. For every intron in CHESS, RefSeq, GENCODE, and MANE, we then computed how many species preserved the consensus dinucleotides (GT and AG) at either end of that intron.

As shown in **Figure 3**, a large majority of introns in protein-coding genes are conserved across all or nearly all 30 species. All four annotation databases show very similar profiles. However, the conservation profile of lncRNAs is quite different from that of protein coding genes, in at least two ways: first, very few introns are conserved across all 30 species, with the largest peak at 20-21 species; and second, the distribution shows a clear secondary peak in introns conserved among 4-7 species. We then computed the most frequent species in which introns from the secondary peak are conserved.

**Figure 3.**
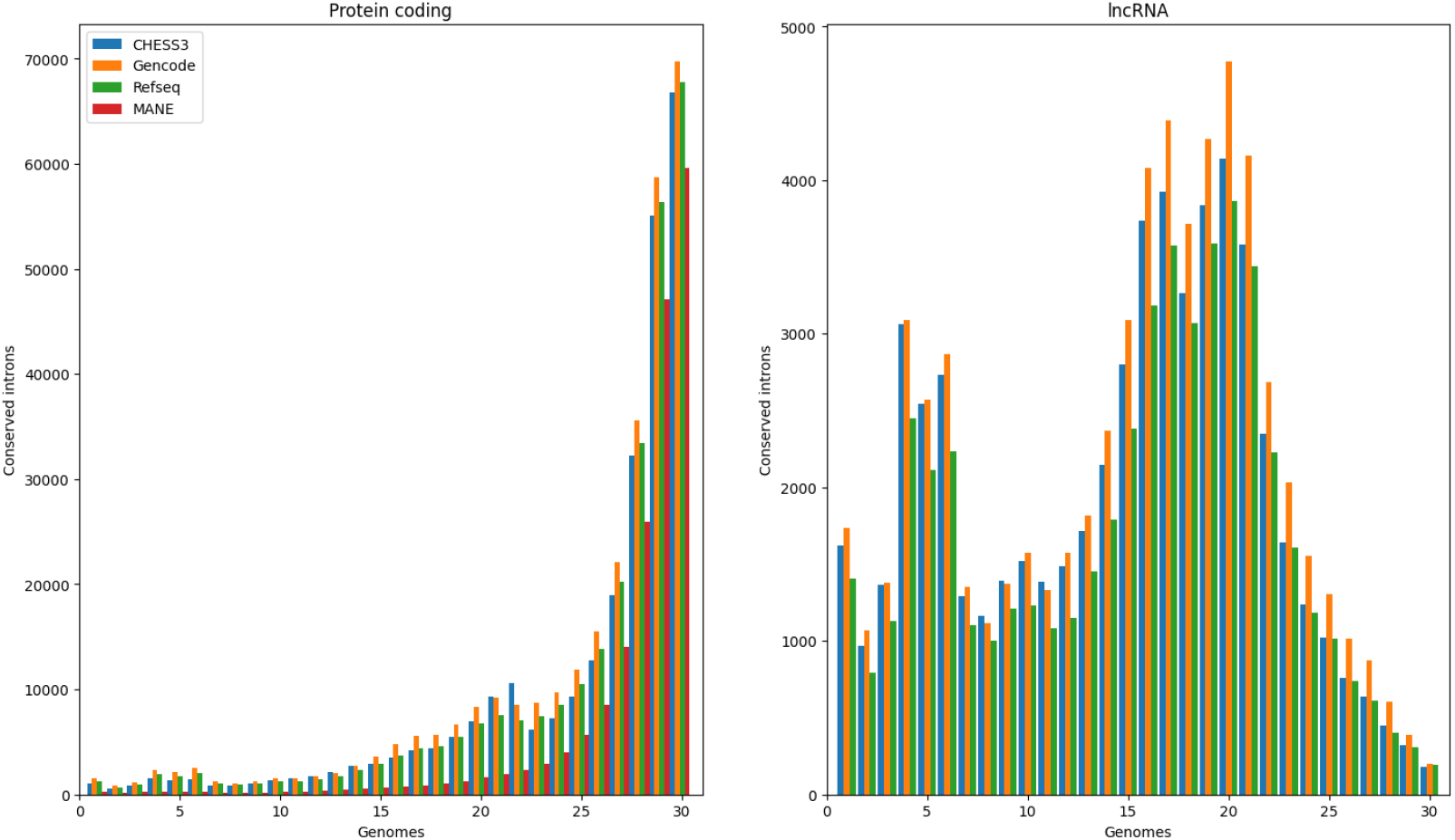
Histogram showing how many introns have both boundaries conserved in a multiple alignment of 27 primates plus 3 additional mammalian species (mouse, dog, and armadillo).

Table 5 shows the ten most frequent such species, where the top five species account for most of the conserved introns. Notably, these lncRNA introns are mostly conserved in apes, with a sharp drop in the number of introns remaining intact outside of this clade. A similar pattern was observed in lncRNAs from the RefSeq and GENCODE databases as well.

**Table 5.** The most common species in which lncRNA introns from the secondary peak of 4-7 species in Figure 3 are conserved. Data shown only for the CHESS database; other annotation databases display highly similar patterns of conservation.

| Species | Number of conserved introns |
| --- | --- |
| Chimpanzee | 7801 |
| Gorilla | 7602 |
| Pygmy chimpanzee (Bonobo) | 7492 |
| Sumatran orangutan | 3676 |
| Northern white-cheeked gibbon | 3517 |
| White-tufted-ear marmoset | 264 |
| Bolivian squirrel monkey | 254 |
| White-faced sapajou | 225 |
| Green monkey | 208 |
| Baboon | 201 |

#### CHESS annotation on CHM13

Although GRCh38 is nearly universally used as the human reference genome, the recently-published CHM13 genome is the first truly complete human genome, adding nearly 200 Mbp of DNA, closing over 900 gaps, and adding thousands of new transcripts, based on the initial annotation [1]. As mentioned above, the annotation of CHM13 was based on GENCODE v35, for which the authors reported 140 new protein-coding genes, and a net increase of 79 protein-coding genes after subtracting genes that were missing in CHM13 (including 23 protein-coding genes that are the result of false duplications in GRCh38).

To produce a more accurate human gene count, and to provide better support for CHM13 as a human reference genome in the future, we mapped all CHESS 3 transcripts onto CHM13 using Liftoff [24], including a routine to find additional gene copies. The resulting annotation, summarized in Supplementary **Table S2**, contains a total of 43,773 genes and 161,410 transcripts, including 2,510 additional gene copies of which 129 are protein coding. In the CHM13 annotation, 19,968 genes are protein coding containing 99,410 transcripts. To summarize the effects of variants in these protein coding genes compared to their GRCh38 sequences, we compared the annotation with LiftoffTools, which identified thousands of synonymous and nonsynonymous changes as well as smaller numbers of frameshifts and indels (**Suppl. Table S3)**.

69 protein coding genes in CHESS 3 failed to map from GRCh38 to CHM 13. Further investigation revealed that all of these genes fell within regions of segmental duplications (typically with >90% identity) in GRCh38, as defined in [25]. This suggests that these gene represent cases where CHM13 has fewer copies of a gene than GRCh38.

#### CHESS annotation on chimpanzee

We also mapped the CHESS 3 annotation onto the chimpanzee (*Pan troglodytes*) genome, Clint_PTRv2 (GenBank accession GCF_002880755.1). A total of 19,343 protein-coding genes mapped over successfully, with an average sequence identity of 98.3% in exons and average coverage of 99.4% of the length of each transcript. A total of 17,227 LncRNA genes mapped to chimpanzee, with an average sequence identity in exons of 96.9% and average coverage of 98.8%. In addition, 95.6% of the introns across all multi-exon genes had perfect agreement on the positions of their splice sites.

## CONCLUSION

The CHESS database uses thousands of RNA sequencing experiments to assemble a comprehensive picture of all human transcripts, each of which has direct experimental evidence of its expression levels. CHESS 3.0 augments this collection with selected, well-annotated genes from the RefSeq, GENCODE, and MANE databases to create a more-complete representation of all genes. The new release of CHESS described here reflects a stricter approach to annotation than in the past, with a greater emphasis on removing transcripts that likely represent non-functional isoforms, and which in turn can hinder downstream analysis when provided to automated genome analysis programs. The result is that CHESS 3.0 has fewer than half as many transcripts as CHESS 2.0, although it has approximately the same number of protein-coding genes. A unique feature of CHESS 3.0 is a complete set of predicted 3D protein structures for >98% of protein-coding genes, which allows users to ask directly how well-ordered these proteins are. Another novel feature is that CHESS 3.0 genes are available on both the older GRCh38 human reference genome and the new, complete CHM13 genome, which contains ∼2,500 more genes. Although the total number of protein-coding genes in CHESS and in other major databases is converging, the number of transcripts remain quite divergent, and much more work is needed before we are likely to have a final picture of all human genes.

## Methods

The pipeline used to create CHESS 3 is presented in **Figure 4**. In summary, 9,795 samples collected across 31 histological types were initially obtained from the GTEx consortium project [26] for the construction of the CHESS catalog version 2.2 [5]. After adding 132 samples that were released by the GTEx consortium in 2018, we aligned the reads with the latest HISAT2 software [27], using an X-only reference genome for female samples to avoid erroneous mapping of reads to the Y chromosome [28], and then assembled aligned reads from the 9,927 samples using StringTie2 [29]. Samples were grouped by tissue type and merged together, as described previously [5]. These initial steps generated 26,335,900 transcripts, the vast majority of which were expressed at low levels.

**Figure 4.**
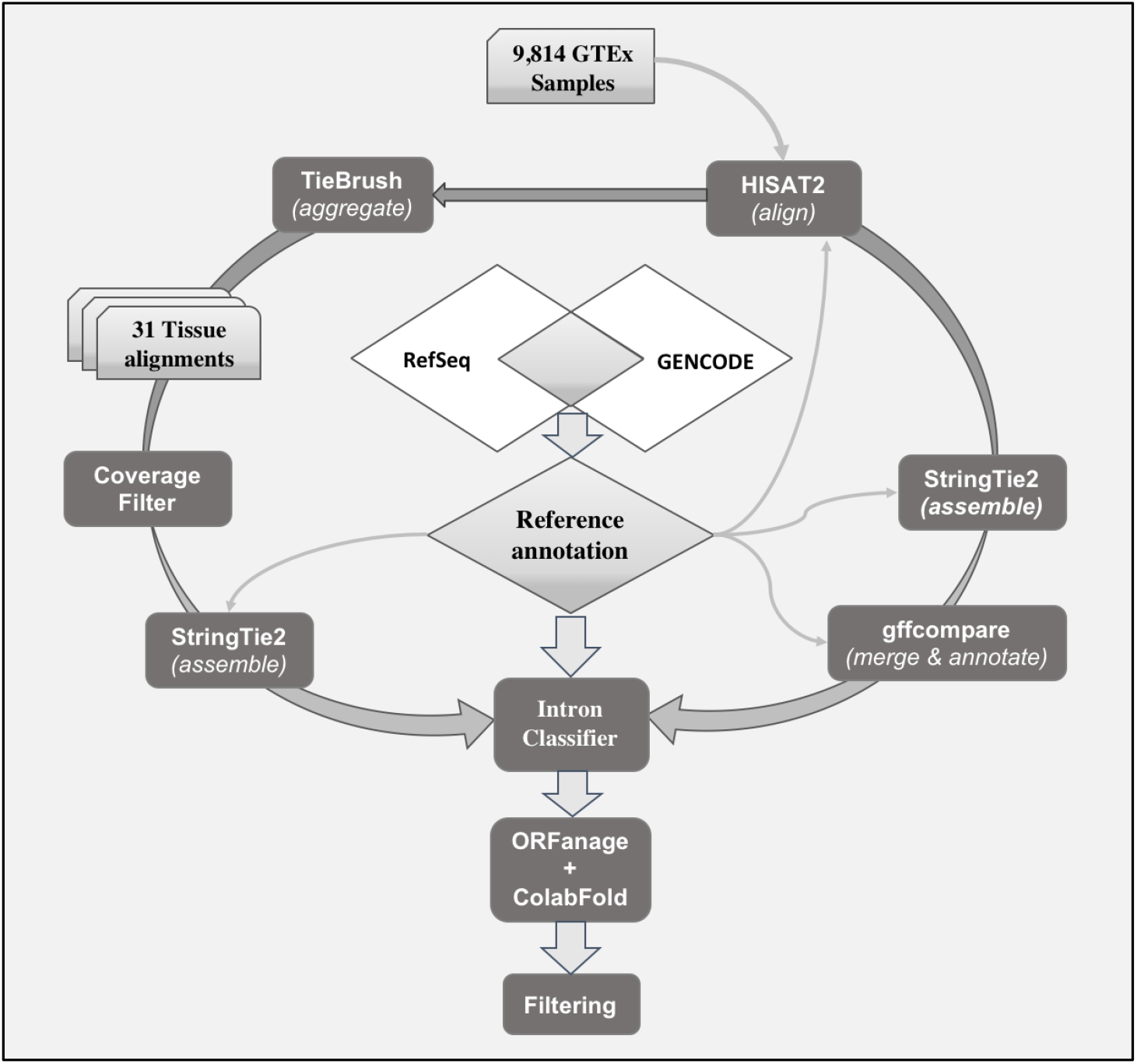
Computational pipeline used to create CHESS 3. First, 9,814 GTEx samples were aligned with HISAT2. Second, the alignments were either directly assembled with StringTie2 or aggregated by tissue with TieBrush. StringTie2’s resulting transcripts were merged and compared to the reference annotation using gffcompare. Low coverage alignments in the “TieBrush”-ed files were filtered out, and the remaining alignments were assembled with StringTie2. Only transcripts that were assembled directly from the individual samples or from “TieBrush”-ed files were retained, and further filtered with an intron classifier designed to recognize introns that resemble most the introns in the reference annotation. ORFanage and ColabFold were used to assign and score ORFs to protein coding transcripts, and pLDDT scores produced by ColabFold were used to filter out low-scoring protein coding transcripts.

We then proceeded through a series of data cleaning and filtering steps, which are described in the Supplementary Methods. These steps were designed to remove transcriptional noise, including transcripts expressed at very low levels as well as fragmented transcripts. To filter out transcripts expressed at very low levels and only in a few samples from a tissue, we aggregated all available alignments from each tissue using TieBrush [30], and reassembled each tissue with StringTie2. We only kept transcripts that were assembled in the initial samples, as well as after aggregating the alignments with TieBrush. We applied further stringent filtering steps to remove noisy transcripts, including only retaining transcripts with well-supported introns. These steps reduced the dataset to 160,482 transcripts, of which 97,661 were protein-coding. All of the protein-coding transcripts were assigned coding sequence (CDS) features either by copying them from matching RefSeq transcripts, where available, or by the ORFanage program as described in Supplementary Methods. For the sake of discussion, we call these the “Beta” proteins here.

We then employed a method not used systematically in previous human gene annotation databases: protein structure prediction by AlphaFold2, which produces highly accurate structures for most proteins [14]. In particular, when the AlphaFold2 pLDDT score is greater than 70, the prediction is considered confident except for short proteins [31].

We began with a less-stringently filtered superset of the assembled transcripts from GTEx, and predicted structures for all proteins shorter than 1000aa using ColabFold [15], a version of AlphaFold2 that runs on public cloud-computing resources, as described in a separate study [32]. This dataset had 194,780 structures. We identified those structures that had pLDDT scores of 70 or above, and we further filtered the set to identify transcripts whose proteins did not match any of the 97,661 “beta” set of proteins. This gave us 54,205 “candidate” transcripts for potential inclusion in CHESS, all of which encoded proteins with scores >= 70 that were not in the Beta set.

We then ran gffcompare to compare the candidate transcripts to the Beta transcripts, and we also ran custom scripts to compare the protein sequences directly. Any proteins that were substrings of the Beta proteins were removed. For proteins that completely contained the Beta proteins (i.e., were longer), we evaluated them based on ColabFold scores: if the ColabFold score was the highest-scoring isoform for a given gene locus, we retained the transcript, otherwise we removed it. These steps reduced the number of candidate transcripts to 31,772. We also noted that if a protein fragment consists largely of well-structured amino acids, it sometimes scores higher than the full-length functional protein, even if it is much shorter. Therefore we removed any predicted proteins that were either (a) shorter than 70aa or (b) less than 2/3 of the length of the longest protein at the same locus. This filtering step reduced the number of candidates to 13,133.

From this set, we removed duplicates in several ways. First, we identified all transcripts that encoded identical proteins at a given gene locus, and if one of the transcripts matched a RefSeq or Gencode transcript, we retained only that one. From the remaining duplicates, we retained the transcript that was assembled in the largest number of GTEx samples. These steps reduced the candidate list to 12,075 transcripts.

Finally, we identified possible conflicting transcripts that overlapped more than one locus, and that might represent read-through transcription. We removed these as well, yielding 11,225 protein-coding transcripts that were then added to the Beta set. Each of these additional transcripts encodes a protein that scored at least 70 and that was not otherwise present in the Beta set.

Annotation of CHM13 used Liftoff [3] to map genes from the primary chromosomes, excluding the alternative scaffolds, onto the complete CHM13 genome. GRCh38 contains a number of regions, mostly on chromosome 21, that are known to be erroneous duplications [33, 34]. These regions contain 15 genes on chr21 that are spurious copies, as well as other spurious genes, and we therefore masked out these genes before mapping the remaining genes onto CHM13. The only exception was TRPM3, which we did not mask out because its erroneous duplications are restricted to intronic regions of GRCh38.

Other than the erroneous duplications, the near-identical ribosomal DNA (rDNA) arrays also present a problem. An rDNA array is composed of several rDNA units, where each unit comprises three ribosomal RNA genes, 18S, 5.8S, and 28S, separated by transcribed spacers and follow by intergenic sequence (IGS) at the end [35]. In CHM13, there are 219 copies of rDNA units located on the acrocentric chromosomes 13, 14, 15, 21 and 22.

We adopted a 2-pass approach to lift over CHESS annotations from GRCh38 to CHM13. First, we masked out all rDNA regions on the CHM13 using bedtools [36] and then mapped all annotations except the rDNA genes onto the masked CHM13 genome, to prevent annotations from being mapped into these complex regions. We used a minimum sequence identity threshold of 95% for identifying additional copies of genes in CHM13. After this initial lift-over process, we merged the rDNA annotations from the CHM13 v2.0 genome into our CHM13 annotations. We used essentially the same Liftoff process (separately) to map the CHESS 3 annotation onto the chimpanzee genome.

## Acknowledgements

This work was supported in part by the U.S. National Institutes of Health grants R01 HG006677, R01 MH123567, and R35 GM130151; by the U.S. National Science Foundation grant DBI-1759518; by National Research Foundation of Korea (NRF) grants 2019R1-A6A1-A10073437, 2020M3-A9G7-103933, 2021-R1C1-C102065, and 2021-M3A9-I4021220]; by Samsung DS research fund; and by the Creative-Pioneering Researchers Program through Seoul National University.

## References

1. Nurk, S., S. Koren, A. Rhie, et al. The complete sequence of a human genome. Science, 2022. 376(6588): p. 44–53.

2. Frankish, A., S. Carbonell-Sala, M. Diekhans, et al. GENCODE: reference annotation for the human and mouse genomes in 2023. Nucleic Acids Res, 2022.

3. Shumate, A. and S.L. Salzberg. Liftoff: accurate mapping of gene annotations. Bioinformatics, 2020. 37(12): p. 1639–43.

4. O’Leary, N.A., M.W. Wright, J.R. Brister, et al. Reference sequence (RefSeq) database at NCBI: current status, taxonomic expansion, and functional annotation. Nucleic Acids Res, 2016. 44(D1): p. D733–45.

5. Pertea, M., A. Shumate, G. Pertea, et al. CHESS: a new human gene catalog curated from thousands of large-scale RNA sequencing experiments reveals extensive transcriptional noise. Genome Biol, 2018. 19(1): p. 208.

6. Hon, C.C., J.A. Ramilowski, J. Harshbarger, et al. An atlas of human long non-coding RNAs with accurate 5’ ends. Nature, 2017. 543(7644): p. 199–204.

7. Rodriguez, J.M., F. Pozo, D. Cerdan-Velez, et al. APPRIS: selecting functionally important isoforms. Nucleic Acids Res, 2022. 50(D1): p. D54–D59.

8. GTEx Consortium. Human genomics. The Genotype-Tissue Expression (GTEx) pilot analysis: multitissue gene regulation in humans. Science, 2015. 348(6235): p. 648–60.

9. Morales, J., S. Pujar, J.E. Loveland, et al. A joint NCBI and EMBL-EBI transcript set for clinical genomics and research. Nature, 2022. 604(7905): p. 310–315.

10. Tress, M.L., F. Abascal, and A. Valencia. Alternative Splicing May Not Be the Key to Proteome Complexity. Trends Biochem Sci, 2017. 42(2): p. 98–110.

11. Varabyou, A., S.L. Salzberg, and M. Pertea. Effects of transcriptional noise on estimates of gene and transcript expression in RNA sequencing experiments. Genome Res, 2020.

12. Kellermayer, D., J.E. Smith, 3rd, and H. Granzier. Novex-3, the tiny titin of muscle. Biophys Rev, 2017. 9(3): p. 201–206.

13. de Morree, A., M. Droog, L. Grand Moursel, et al. Self-regulated alternative splicing at the AHNAK locus. FASEB J, 2012. 26(1): p. 93–103.

14. Jumper, J., R. Evans, A. Pritzel, et al. Highly accurate protein structure prediction with AlphaFold. Nature, 2021. 596(7873): p. 583–589.

15. Mirdita, M., K. Schutze, Y. Moriwaki, et al. ColabFold: making protein folding accessible to all. Nat Methods, 2022. 19(6): p. 679–682.

16. Tunyasuvunakool, K., J. Adler, Z. Wu, et al. Highly accurate protein structure prediction for the human proteome. Nature, 2021. 596(7873): p. 590–596.

17. Zhao, L., J. Wang, Y. Li, et al. NONCODEV6: an updated database dedicated to long non-coding RNA annotation in both animals and plants. Nucleic Acids Res, 2021. 49(D1): p. D165–D171.

18. Volders, P.J., J. Anckaert, K. Verheggen, et al. LNCipedia 5: towards a reference set of human long non-coding RNAs. Nucleic Acids Res, 2019. 47(D1): p. D135–D139.

19. Seifuddin, F., K. Singh, A. Suresh, et al. lncRNAKB, a knowledgebase of tissue-specific functional annotation and trait association of long noncoding RNA. Sci Data, 2020. 7(1): p. 326.

20. RNAcentral Consortium. RNAcentral 2021: secondary structure integration, improved sequence search and new member databases. Nucleic Acids Res, 2021. 49(D1): p. D212–D220.

21. Pertea, M., A. Shumate, G. Pertea, et al. Thousands of large-scale RNA sequencing experiments yield a comprehensive new human gene list and reveal extensive transcriptional noise. bioRxiv, 2018.

22. Ponting, C.P. and W. Haerty. Genome-Wide Analysis of Human Long Noncoding RNAs: A Provocative Review. Annu Rev Genomics Hum Genet, 2022. 23: p. 153–172.

23. Kent, W.J., C.W. Sugnet, T.S. Furey, et al. The human genome browser at UCSC. Genome Res, 2002. 12(6): p. 996–1006.

24. Shumate, A. and S.L. Salzberg. Liftoff: accurate mapping of gene annotations. Bioinformatics, 2020.

25. Alkan, C., J.M. Kidd, T. Marques-Bonet, et al. Personalized copy number and segmental duplication maps using next-generation sequencing. Nat Genet, 2009. 41(10): p. 1061–7.

26. GTEx Consortium. The Genotype-Tissue Expression (GTEx) project. Nat Genet, 2013. 45(6): p. 580–5.

27. Kim, D., J.M. Paggi, C. Park, et al. Graph-based genome alignment and genotyping with HISAT2 and HISAT-genotype. Nature Biotechnology, 2019. 37(8): p. 907–915.

28. Olney, K.C., S.M. Brotman, J.P. Andrews, et al. Reference genome and transcriptome informed by the sex chromosome complement of the sample increase ability to detect sex differences in gene expression from RNA-Seq data. Biol Sex Differ, 2020. 11(1): p. 42.

29. Kovaka, S., A.V. Zimin, G.M. Pertea, et al. Transcriptome assembly from long-read RNA-seq alignments with StringTie2. Genome Biol, 2019. 20(1): p. 278.

30. Varabyou, A., G. Pertea, C. Pockrandt, and M. Pertea. TieBrush: an efficient method for aggregating and summarizing mapped reads across large datasets. Bioinformatics, 2021. 37(20): p. 3650–3651.

31. Monzon, V., D.H. Haft, and A. Bateman. Folding the unfoldable: using AlphaFold to explore spurious proteins. Bioinformatics Advances, 2022. 2(1).

32. Sommer, M.J., S. Cha, A. Varabyou, et al. Structure-guided isoform identification for the human transcriptome. Elife, 2022. 11.

33. Aganezov, S., S.M. Yan, D.C. Soto, et al. A complete reference genome improves analysis of human genetic variation. Science, 2022. 376(6588): p. eabl3533.

34. Miller, C.A., J.R. Walker, T.L. Jensen, et al. Failure to Detect Mutations in U2AF1 due to Changes in the GRCh38 Reference Sequence. J Mol Diagn, 2022. 24(3): p. 219–223.

35. Agrawal, S. and A.R.D. Ganley. The conservation landscape of the human ribosomal RNA gene repeats. PLoS One, 2018. 13(12): p. e0207531.

36. Quinlan, A.R. and I.M. Hall. BEDTools: a flexible suite of utilities for comparing genomic features. Bioinformatics, 2010. 26(6): p. 841–2.

