## Supplemental Materials and Methods for "CHESS 3: an improved, comprehensive catalog of human genes and transcripts based on large-scale expression data, phylogenetic analysis, and protein structure"

### Supplementary Methods

#### Reference genome sequences and annotation

In the GRCh38 human genome [1], numerous sequences are included as patches to the main assembly, as well as alternative forms of specific regions along the genome. Because the variations in these sequences are typically minor, many of the genetic sequences in these "alt" scaffolds appear twice (or more) in the GRCh38 assembly, even when they occur just once in the genome. These artificial duplications can confuse alignment programs, causing them to report that a read is multi-mappe or even unmapped in some cases. To avoid these problems, for all our analyses we relied exclusively on a non-redundant set of sequences from GRCh38 patch 12 (GenBank accession GCA_000001405.27). These sequences included all available primary chromosome scaffolds (1-22, X, Y, and M) plus the set of unplaced contigs. Similarly, we removed alternative scaffolds and patch sequences from the RefSeq v110 [2], GENCODE v31 [3], and CHESS v2.2 [4] annotations which were used in several stages of our analysis.

The sex chromosomes in humans present a unique set of challenges for annotation. The X and Y chromosomes in humans contain several large regions of near-perfect identity, typically referred to as pseudo-autosomal regions (PARs). Mapping reads from a female sample to an assembly that includes both X and Y will inevitably result in some reads from PARs being mapped erroneously to the PARs on Y chromosome, thus leading to a decreased abundance for genes on the X chromosome [5]. To address this problem, we aligned all samples designated as female to a copy of GRCh38 with the Y chromosome removed.

To create the version of CHESS 3 mapped to the complete, gap-free human genome, we used the recently published CHM13 genome [6], augmented with the finished sequence of chromosome Y from another individual (because CHM13 was a female sample), which is designed CHM13 2.0 (GenBank accession GCA_009914755.4).

The choice of annotation to be used by mapping and assembly software to guide the analysis is an important consideration as well. The RefSeq and GENCODE human annotation databases disagree on a majority of their gene isoforms, although they roughly agree on the locations of the protein-coding genes [4, 7]. Including poorly supported exon-intron structures in our analysis would inevitably bias both the mapping process and splice junction inference, which in turn would bias the assembly of transcripts in favor of potentially erroneous features. On the other hand, a completely de novo approach to the alignment and assembly problems (i.e., not relying on annotation) decreases the accuracy of spliced alignment [8, 9].

To tackle this problem, we created a conservative annotation database to guide both spliced alignment and assembly of RNA-seq reads. We chose RefSeq and GENCODE as the two most widely-used human gene catalogs, and extracted a subset of the transcripts for which both catalogs had precisely the same introns annotated. We then used this intersection of the two databases to guide alignment.

**Alignment and Assembly**

Samples were mapped to the sex-specific versions of GRCh38 with HISAT2 version 2.2 [9]. HISAT2 indices for each sex were augmented with the donor-acceptor site information from the intersection of RefSeq and GENCODE described above. For this analysis we used a version of HISAT2 that includes an "--rna-sensitive" flag. This flag guarantees that for all multi-mapping reads at least one mapping will be reported, unlike the default behavior of HISAT2, which does not report mappings for reads that align to more than 10 locations on the genome. These changes led to a noticeable increase in the proportions of aligned reads as compared to the alignment methods used when building the original CHESS database, as shown in Supplementary **Figure S1**.

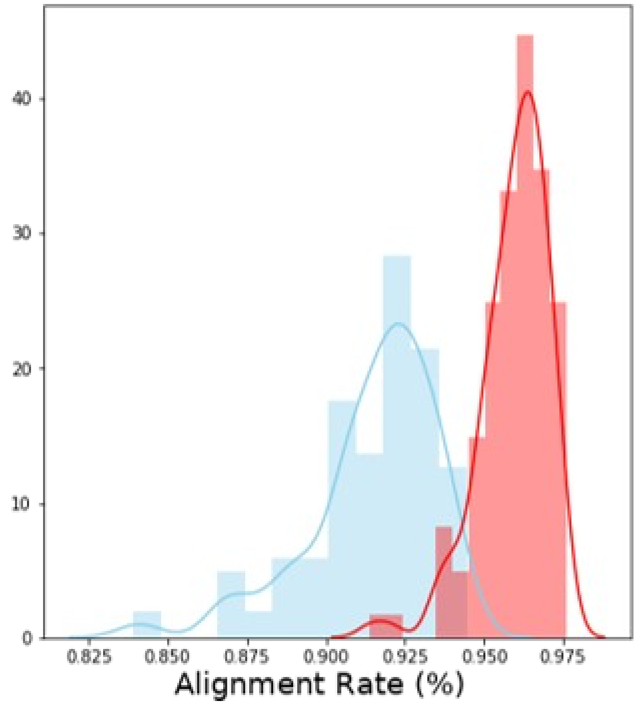


**Supplementary Figure S1.** Change in the alignment rate of the GTEx samples between the alignment strategy implemented in CHESS2 (blue) and the improved alignment protocol using an updated HISAT2 release, sex-specific indices, and newer guide annotation (red).

Following the protocol used for the previous releases of CHESS, we discarded 133 samples having an anomalously low total number of reads or mapping rate. These samples were identified using the interquartile range (IQR) method with a scaling factor of 1.5 following standard practices (Supplemental **Table S4**).

Mapped reads were then assembled using StringTie 2 [10] in its reference-guided mode with default parameters. Isoforms with identical intron chains from assembled samples were merged using gffcompare [11] for each tissue type as defined in the GTEx metadata. Finally, merged results from the tissues were merged once more to obtain the final set of unique transcripts across the entire dataset.

##### Quantification of properties of splice junctions

In our evaluations of the previous version of CHESS, we hypothesized that many apparently novel introns were mapping artifacts. While much effort in our re-evaluation was dedicated to the refinement of mapping and assembly through tuning of parameters and a refined reference annotation, some of the novel splice junctions identified by HISAT2 might still be artifacts of the spliced-alignment algorithm or errors in the read data. To select the best candidates for novel isoforms, we first focused our attention on selecting the best candidate introns based on observations from read mappings.

From a computational perspective, an intron is a well-defined quantifiable unit of gene expression, which has motivated many studies that collected and quantified splicing events rather than trying to assemble full transcripts [12, 13]. Unlike exons, which are described by a range of coordinates and have natural and artifactual perturbations in coverage, splice junctions can be identified by two positions that are spanned by a single read.

We began our analysis of splice junctions from alignments by collapsing alignments of all samples for each tissue type separately. We chose to process data for each of the 31 tissue types independently due to expected and previously reported homogeneity of expression within tissues [14]. In this step, for each intron in each tissue we recorded:

1) the number of samples in which the intron was observed,
2) the coverage of the exon-exon junction by uniquely mapped reads,
3) the ratio of coverage by uniquely mapped reads to the total number of mapped reads that cover the junction,
4) the ratio of coverage at the donor site position to the coverage at the first intronic position,
5) the ratio of coverage at the acceptor site position to the coverage at the last intronic position,
6) the maximum number of bases by which a single read extends upstream and downstream of the donor and acceptor sites respectively,
7) the number of assembled transcripts in which the junction was observed,
8) the ratio of coverage by uniquely mapped reads between the positive and negative strands, and
9) whether or not the junction was present in the guide annotation.

##### Training a classifier for intron junctions

Having collected information about the splice junctions from the aligner, we extracted a subset of junctions that were present in the conservative guide annotation. Although genes in the guide annotation are likely to be genuine, certain cell-type-specific genes and isoforms might not be actively expressed in each of the groups being analyzed. Low traces of transcripts from these regions might still be detected due to noisy transcription by RNA polymerase II or contamination during sample preparation [12-16]. To filter such molecules out of our training set, we treated any known splice junction as novel if its total coverage in a tissue by uniquely mapped reads fell below 100.

For each tissue type, we randomly chose 80% of the novel and known junctions for training a machine learning model, and reserved 20% for validation. We used LightGBM, a random forest decision tree method [17], and fit the model to the training data to obtain a decision boundary that best separated known junctions from novel ones. The model parameters were tuned to account for the imbalance of binary labels in the training data. The model was trained for 50,000 rounds or until no further improvement was observed for 500 consecutive rounds with a learning rate of 0.005. Other parameters, such as the number of leaf nodes, maximum depth, minimum number of datapoints in leaves, bagging fraction, and fraction of features to be used for each round of boosting were selected via an exhaustive grid search.

The model was then used to predict labels for all splice junctions in each validation set. All splice junctions labeled as valid by the model were reserved for the downstream analysis.

**Transcriptional and technical noise reduction using TieBrush**

To improve the specificity of the downstream analysis, we set out to reduce transcriptional and technical noise at the level of alignment and assembly. We used TieBrush [18] to aggregate all available alignments from each tissue in the GTEx dataset. The resulting 31 aggregated alignment files were filtered based on coverage, retaining only those reads spanning positions with coverage greater than 3 across all samples in the collapsed representation of a tissue. We used StringTie2 to assemble the filtered alignment files, followed by gffcompare to merge all transcripts assembled from the 31 tissues. Only 987,244 transcripts remained from the originally assembled ~26.3 million transcripts in all individual samples after filtering with TieBrush, and we retained these as potential candidates for our future analyses, henceforward referred to as the “union set" of transcripts.

**Selection of transcripts novel to the CHESS catalog**

We then processed transcripts from the union set as follows. For each tissue type, we computed the total and average TPM values and the number of samples in which each assembled isoform was seen. Similar to the previous CHESS build, we imposed thresholds on the minimum number of samples and average TPM threshold of 10 and 1 respectively. Because the selection was done in a tissue-specific manner in this CHESS build, compared to aggregating results across all of the GTEx dataset, the thresholds were more restrictive. We also excluded transcripts with expression levels that were less than 10% of the highest expressed transcript at a locus.

Having a set of splice junctions predicted as valid from the previous step, we developed a custom algorithm to select a set of isoforms that would explain all of the valid introns. First, we sorted all transcripts in descending order by cumulative TPM computed over all samples in each group. Iterating from most abundant to least abundant isoform, our algorithm did the following:

1. Sort all transcripts in descending order by cumulative TPM computed over all samples in each group.
2. For each transcript *T* (most abundant to least abundant), if *T* contains valid introns that have not been seen in more-abundant transcripts, then:
    a. Add *T* to the set of transcripts to retain, and
    b. remove all of *T*'s introns from the list of available valid introns.

At the end of the run, the algorithm produced a concise set of isoforms that covered all splice junctions labeled as valid by the machine learning model.

**Adding novel single-exon transcripts**

Thus far our analysis relied on utilizing properties of splice junctions to identify novel transcripts to be included in CHESS, thus excluding single-exon transcripts. To re-introduce high quality single-exon transcripts into the catalog, we screened each tissue for single-exon transcripts assembled in the same minimum number of samples as required for the multi-exon transcripts. Furthermore, we kept single-exon transcripts only if they had mean expression above the threshold computed for each tissue using the IQR method with a scaling factor of 1.5, per standard practice.

**Additional filtering and adjustments**

Because the 3’ and 5’ ends of transcripts are hard to determine precisely from RNA-Seq data, we adjusted the 3’ and 5’ ends of the assembled transcripts to the nearest 3’ and 5’ ends of transcripts in either RefSeq or GENCODE present in the same locus, ensuring that ORFs, if available, were preserved.

Second, we replaced all transcripts on the mitochondrial DNA, for which annotation is complete and well-established, with the RefSeq annotation, rather than using transcripts assembled by StringTie.

Next, we removed all readthrough transcripts between loci, unless they were also present in the RefSeq Select annotation. We define a readthrough transcript as any transcript that overlaps multiple loci on the same strand after the 3’ and 5’ end correction.

**Adding transcripts from known sources**

We added transcripts from known sources based on the following criteria. First, we decided to include all transcripts in the recently-published MANE database [7] a joint effort of NCBI and EMBL-EBI to create a database with just one very high-quality transcript for each protein-coding gene. The vast majority of these were already in CHESS after the assembly steps above, but we identified and added a small number of MANE transcripts to ensure completeness.

We also added several categories of transcripts documented in RefSeq and GENCODE, namely: (1) any transcripts GENCODE that were assembled in at least one GTEx sample and that were present either in RefSeq Select or in both RefSeq and GENCODE; (2) RefSeq transcripts representing VDJ segments and Y RNA; (3) RefSeq transcripts for tRNAs, tmRNAs, rRNAs, and nucleolar RNAs.

**Annotation of Open Reading Frames**

All protein-coding transcripts in CHESS have coding sequence (CDS) features that specify the boundaries of the open reading frame (ORF). For protein-coding transcripts that appear either in MANE, RefSeq, or GENCODE we copied the ORF boundaries over to the corresponding CHESS transcript, preferring the RefSeq ORF annotation if the transcript was both in RefSeq and GENCODE. To identify boundaries of ORFs in novel protein-coding transcripts, we used ORFanage [19] with default parameters, guided by the annotation of full-length proteins from RefSeq. We made sure sure to preserve seleno-proteins and alternative (i.e., non-ATG) start codons where annotated. Finally, we used the ORFs of the the 224 protein-coding genes that were novel in CHESS 2 [4], to annotate potential-coding regions of transcripts that were in non-protein coding loci in both RefSeq and GENCODE .

#### References


1. Schneider, V.A., T. Graves-Lindsay, K. Howe, N. Bouk, H.C. Chen, P.A. Kitts, . . . D.M. Church, Evaluation of GRCh38 and de novo haploid genome assemblies demonstrates the enduring quality of the reference assembly. *Genome Res*, 2017. **27**(5): p. 849-864.
2. O'Leary, N.A., M.W. Wright, J.R. Brister, S. Ciufo, D. Haddad, R. McVeigh, . . . K.D. Pruitt, Reference sequence (RefSeq) database at NCBI: current status, taxonomic expansion, and functional annotation. *Nucleic Acids Res*, 2016. **44**(D1): p. D733-45.
3. Frankish, A., M. Diekhans, A.M. Ferreira, R. Johnson, I. Jungreis, J. Loveland, . . . P. Flicek, GENCODE reference annotation for the human and mouse genomes. *Nucleic Acids Res*, 2019. **47**(D1): p. D766-D773.
4. Pertea, M., A. Shumate, G. Pertea, A. Varabyou, F.P. Breitwieser, Y.C. Chang, . . . S.L. Salzberg, CHESS: a new human gene catalog curated from thousands of large-scale RNA sequencing experiments reveals extensive transcriptional noise. *Genome Biol*, 2018. **19**(1): p. 208.
5. Olney, K.C., S.M. Brotman, J.P. Andrews, V.A. Valverde-Vesling, and M.A. Wilson, Reference genome and transcriptome informed by the sex chromosome complement of the sample increase ability to detect sex differences in gene expression from RNA-Seq data. *Biol Sex Differ*, 2020. **11**(1): p. 42.

6. Nurk, S., S. Koren, A. Rhie, M. Rautiainen, A.V. Bzikadze, A. Mikheenko, . . . A.M. Phillippy, The complete sequence of a human genome. *Science*, 2022. **376**(6588): p. 44-53.
7. Morales, J., S. Pujar, J.E. Loveland, A. Astashyn, R. Bennett, A. Berry, . . . T.D. Murphy, A joint NCBI and EMBL-EBI transcript set for clinical genomics and research. *Nature*, 2022. **604**(7905): p. 310-315.
8. Dobin, A., C.A. Davis, F. Schlesinger, J. Drenkow, C. Zaleski, S. Jha, . . . T.R. Gingeras, STAR: ultrafast universal RNA-seq aligner. *Bioinformatics*, 2013. **29**(1): p. 15-21.
9. Kim, D., J.M. Paggi, C. Park, C. Bennett, and S.L. Salzberg, Graph-based genome alignment and genotyping with HISAT2 and HISAT-genotype. *Nat Biotechnol*, 2019. **37**(8): p. 907-915.
10. Kovaka, S., A.V. Zimin, G.M. Pertea, R. Razaghi, S.L. Salzberg, and M. Pertea, Transcriptome assembly from long-read RNA-seq alignments with StringTie2. *Genome Biol*, 2019. **20**(1): p. 278.
11. Pertea, G. and M. Pertea, GFF Utilities: GffRead and GffCompare. *F1000Research*, 2020. **9**(304).
12. Collado-Torres, L., A. Nellore, K. Kammers, S.E. Ellis, M.A. Taub, K.D. Hansen, . . . J.T. Leek, Reproducible RNA-seq analysis using recount2. *Nat Biotechnol*, 2017. **35**(4): p. 319-321.
13. Li, Y.I., D.A. Knowles, J. Humphrey, A.N. Barbeira, S.P. Dickinson, H.K. Im, and J.K. Pritchard, Annotation-free quantification of RNA splicing using LeafCutter. *Nat Genet*, 2018. **50**(1): p. 151-158.
14. Varabyou, A., S.L. Salzberg, and M. Pertea, Effects of transcriptional noise on estimates of gene and transcript expression in RNA sequencing experiments. *Genome Res*, 2021. **31**: p. 301-308.
15. Struhl, K., Transcriptional noise and the fidelity of initiation by RNA polymerase II. *Nat Struct Mol Biol*, 2007. **14**(2): p. 103-5.
16. Cavallaro, M., M.D. Walsh, M. Jones, J. Teahan, S. Tiberi, B. Finkenstadt, and D. Hebenstreit, 3 (')-5 (') crosstalk contributes to transcriptional bursting. *Genome Biol*, 2021. **22**(1): p. 56.
17. Ke, G., Q. Meng, T. Finley, T. Wang, W. Chen, W. Ma, . . . T.-Y. Liu. *LightGBM: A Highly Efficient Gradient Boosting Decision Tree*. in *Advances in Neural Information Processing Systems*. 2017.
18. Varabyou, A., G. Pertea, C. Pockrandt, and M. Pertea, TieBrush: an efficient method for aggregating and summarizing mapped reads across large datasets. *Bioinformatics*, 2021. **37**(20): p. 3650–3651.
19. Varabyou, A., ORFanage (2021), GitHub repository, https://github.com/alevar/ORFanage.

#### Supplementary Tables

**Table S1. Novel, highly-expressed genes in CHESS where at least one transcript encodes a protein sequence that is not found in either RefSeq or GENCODE. Each gene has a cumulative TPM >1000 across all GTEx samples, and the novel proteins account for >50% of the total expression across all samples. The column labelled MANE Gene ID shows the corresponding identifier from the MANE database.**

| CHESS Gene ID | MANE Gene ID | Gene Name | Gene TPM (SUM) | Novel Expression (%) |
|---|---|---|---|---|
| CHS.31993 | ENSG00000197756.10 | RPL37A | 15948994.3 | 0.90309901 |
| CHS.8031 | ENSG00000166441.13 | RPL27A | 12881838.8 | 0.74281356 |
| CHS.8948 | ENSG00000254772.1 | EEF1G | 12728611.3 | 0.6985990 |

| | | | | |
|---|---|---|---|---|
| | 0 | | | 6 |
| CHS.42691 | ENSG00000145592.14 | RPL37 | 5555090.1 | 0.85229835 |
| CHS.53998 | ENSG00000156467.10 | UQCRB | 2111561.67 | 0.77249169 |
| CHS.21644 | ENSG00000109046.15 | WSB1 | 1367507.68 | 0.87281611 |
| CHS.29912 | ENSG00000115307.17 | AUP1 | 1244619.81 | 0.54114632 |
| CHS.2468 | ENSG00000213366.13 | GSTM2 | 1190872.61 | 0.59886567 |
| CHS.14651 | ENSG00000134884.15 | ARGLU1 | 1069204.14 | 0.90077677 |
| CHS.18243 | ENSG00000182768.9 | NGRN | 1033147.3 | 0.95718703 |
| CHS.10637 | ENSG00000002016.18 | RAD52 | 850262.45 | 0.93342821 |
| CHS.8911 | ENSG00000221968.9 | FADS3 | 831407.76 | 0.52590915 |
| CHS.20156 | ENSG00000102901.13 | CENPT | 625406.98 | 0.95496481 |
| CHS.18822 | ENSG00000167969.13 | ECI1 | 594796.15 | 0.82833046 |
| CHS.25677 | ENSG00000213339.9 | QTRT1 | 570159.94 | 0.5887341 |
| CHS.4860 | ENSG00000135744.9 | AGT | 563459.24 | 0.58930445 |
| CHS.25465 | ENSG00000088247.19 | KHSRP | 559635.44 | 0.55557218 |
| CHS.7054 | ENSG00000059915.17 | PSD | 550258.36 | 0.80345667 |
| CHS.35908 | ENSG00000100359.21 | SGSM3 | 527315.56 | 0.58567451 |
| CHS.11730 | ENSG00000094914.14 | AAAS | 464197.28 | 0.52852944 |
| CHS.9230 | ENSG00000173992.9 | CCS | 438105.77 | 0.51805446 |
| CHS.34209 | ENSG00000125520.14 | SLC2A4RG | 435611.42 | 0.58943058 |
| CHS.9443 | ENSG00000214530.10 | STARD10 | 430676.44 | 0.66061638 |
| CHS.6055 | ENSG00000107551.21 | RASSF4 | 426034.17 | 0.52350146 |
| CHS.17863 | ENSG00000140474.1 | ULK3 | 400730.94 | 0.7151924 |

| | | | | |
|---|---|---|---|---|
| | 4 | | | 9 |
| CHS.53339 | ENSG00000164808.17 | SPIDR | 382635.74 | 0.51772919 |
| CHS.16973 | ENSG00000215252.12 | GOLGA8B | 365944.57 | 0.97109338 |
| CHS.8635 | ENSG00000109920.13 | FNBP4 | 337876.69 | 0.60120939 |
| CHS.10726 | ENSG00000111254.8 | AKAP3 | 327272.87 | 0.91989965 |
| CHS.54817 | ENSG00000182307.14 | C8orf33 | 310117.14 | 0.52524191 |
| CHS.47261 | ENSG00000196591.12 | HDAC2 | 286602.82 | 0.71685659 |
| CHS.20122 | ENSG00000196123.13 | KIAA0895L | 279343.97 | 0.63038884 |
| CHS.888 | ENSG00000158062.21 | UBXN11 | 265034.2 | 0.62351591 |
| CHS.24271 | ENSG00000167088.11 | SNRPD1 | 258924.22 | 0.72113767 |
| CHS.33307 | ENSG00000171456.21 | ASXL1 | 255796.06 | 0.5123238 |
| CHS.38393 | ENSG00000240303.8 | ACAD11 | 255382.53 | 0.78432812 |
| CHS.40975 | ENSG00000187758.8 | ADH1A | 240972.08 | 0.75749282 |
| CHS.13349 | ENSG00000112787.14 | FBRSL1 | 238768.74 | 0.54048708 |
| CHS.5374 | ENSG00000187134.14 | AKR1C1 | 235809.87 | 0.5736636 |
| CHS.26064 | ENSG00000105647.19 | PIK3R2 | 222610.48 | 0.52743784 |
| CHS.18658 | ENSG00000076344.16 | RGS11 | 220705.6 | 0.63292631 |
| CHS.21604 | ENSG00000178307.10 | TMEM11 | 206453.22 | 0.57868625 |
| CHS.47128 | ENSG00000146285.14 | SCML4 | 205173.06 | 0.9828727 |
| CHS.40538 | ENSG00000185873.8 | TMPRSS11B | 197752.6 | 0.53145127 |
| CHS.46625 | ENSG00000151914.22 | DST | 192935.92 | 0.58914447 |
| CHS.56137 | ENSG00000148120.18 | AOPEP | 187310.48 | 0.83215178 |

| | | | | |
|---|---|---|---|---|
| CHS.22035 | ENSG00000273559.5 | CWC25 | 186978.21 | 0.50994354 |
| CHS.25841 | ENSG00000037757.14 | MRI1 | 185856.41 | 0.67510424 |
| CHS.35807 | ENSG00000184381.20 | PLA2G6 | 184800.18 | 0.57866248 |
| CHS.44852 | ENSG00000160883.11 | HK3 | 180510.53 | 0.63601137 |
| CHS.45529 | ENSG00000137177.20 | KIF13A | 174377.35 | 0.60214506 |
| CHS.23510 | ENSG00000167302.11 | TEPSIN | 168267.31 | 0.56992407 |
| CHS.38207 | ENSG00000173706.14 | HEG1 | 150366.26 | 0.82126456 |
| CHS.2833 | ENSG00000198483.13 | ANKRD35 | 148020.61 | 0.56842341 |
| CHS.15633 | ENSG00000126790.12 | L3HYPDH | 144260.55 | 0.54442826 |
| CHS.4452 | ENSG00000082512.15 | TRAF5 | 138951.22 | 0.57141808 |
| CHS.11884 | ENSG00000135473.16 | PAN2 | 136201.52 | 0.63878443 |
| CHS.2374 | ENSG00000237763.10 | AMY1A | 131596.44 | 0.95900482 |
| CHS.41695 | ENSG00000109756.10 | RAPGEF2 | 129034.72 | 0.65526038 |
| CHS.27408 | ENSG00000182310.15 | SPACA6 | 120630.91 | 0.70187285 |
| CHS.8523 | ENSG00000110455.14 | ACCS | 120212.51 | 0.69970097 |
| CHS.16455 | ENSG00000156381.9 | ANKRD9 | 117445.72 | 0.71945644 |
| CHS.150 | ENSG00000149527.18 | PLCH2 | 115646.57 | 0.50073824 |
| CHS.58993 | ENSG00000102287.19 | GABRE | 115563.64 | 0.96460401 |
| CHS.45909 | ENSG00000137337.16 | MDC1 | 115004.83 | 0.5233155 |
| CHS.13118 | ENSG00000196498.14 | NCOR2 | 114605.68 | 0.9594706 |
| CHS.7787 | ENSG00000149043.17 | SYT8 | 113638.47 | 0.55899793 |
| CHS.18712 | ENSG00000127586.1 | CHTF18 | 105188.67 | 0.6086655 |

| | | | | |
|---|---|---|---|---|
| | 7 | | | 5 |
| CHS.11631 | ENSG00000139610.2 | CELA1 | 103686.47 | 0.99641699 |
| CHS.34153 | ENSG00000101194.18 | SLC17A9 | 101041.97 | 0.74185272 |
| CHS.3509 | ENSG00000162755.14 | KLHDC9 | 100400.74 | 0.59294135 |
| CHS.2077 | ENSG00000162642.14 | C1orf52 | 98656.55 | 0.637828 |
| CHS.23235 | ENSG00000170190.16 | SLC16A5 | 96356.69 | 0.53327434 |
| CHS.25172 | ENSG00000064687.13 | ABCA7 | 95168.14 | 0.74546103 |
| CHS.5268 | ENSG00000047056.17 | WDR37 | 93245.26 | 0.5169895 |
| CHS.50362 | ENSG00000011426.11 | ANLN | 93146.11 | 0.66215293 |
| CHS.37523 | ENSG00000189283.10 | FHIT | 88806.11 | 1 |
| CHS.31479 | ENSG00000155657.29 | TTN | 87338.28 | 0.62397943 |
| CHS.36048 | ENSG00000138964.17 | PARVG | 83913.9 | 0.54302684 |
| CHS.12793 | ENSG00000173064.14 | HECTD4 | 81951.68 | 0.86851179 |
| CHS.46564 | ENSG00000096093.16 | EFHC1 | 77849 | 0.79268854 |
| CHS.10624 | ENSG00000139044.12 | B4GALNT3 | 77780.45 | 0.54082356 |
| CHS.51771 | ENSG00000106344.8 | RBM28 | 77213.23 | 0.50483045 |
| CHS.371 | ENSG00000009724.18 | MASP2 | 76686.13 | 0.64257878 |
| CHS.7781 | ENSG00000244242.2 | IFITM10 | 75508.54 | 0.8015153 |
| CHS.12022 | ENSG00000177990.12 | DPY19L2 | 72758.15 | 0.6409462 |
| CHS.37579 | ENSG00000151276.24 | MAGI1 | 72075.78 | 0.70737354 |
| CHS.36260 | ENSG00000177989.14 | ODF3B | 70359.52 | 0.58870697 |
| CHS.26991 | ENSG00000283632.4 | EXOC3L2 | 67597.05 | 0.96716765 |
| CHS.49886 | ENSG00000106415.1 | GLCCI1 | 64306.5 | 0.5017194 |

| | | | | |
|---|---|---|---|---|
| | 3 | | | 2 |
| CHS.2376 | ENSG00000174876.17 | AMY1B | 64027.42 | 0.69079607 |
| CHS.10856 | ENSG00000173262.12 | SLC2A14 | 61341.46 | 0.87156305 |
| CHS.28885 | ENSG00000183891.6 | TTC32 | 59665.87 | 0.68030785 |
| CHS.35379 | ENSG00000133460.20 | SLC2A11 | 57896.76 | 0.50471443 |
| CHS.40709 | ENSG00000138771.16 | SHROOM3 | 56344.07 | 0.75778463 |
| CHS.46063 | ENSG00000237541.4 | HLA-DQA2 | 52629.03 | 0.72421057 |
| CHS.4794 | ENSG00000154358.23 | OBSCN | 52184.16 | 0.80375712 |
| CHS.17417 | ENSG00000181827.16 | RFX7 | 51966.04 | 1 |
| CHS.43096 | ENSG00000249437.8 | NAIP | 50223.05 | 0.85191799 |
| CHS.52219 | ENSG00000284691.1 | ENSG00000284691 | 48038.77 | 0.50476709 |
| CHS.17326 | ENSG00000140287.11 | HDC | 46734.45 | 0.80059421 |
| CHS.48169 | ENSG00000184786.6 | DYNLT2 | 45863.05 | 0.89485392 |
| CHS.47327 | ENSG00000111877.18 | MCM9 | 44725.52 | 0.55301112 |
| CHS.29769 | ENSG00000115977.21 | AAK1 | 42285.76 | 0.65867327 |
| CHS.51973 | ENSG00000146858.8 | ZC3HAV1L | 41873.86 | 0.90042571 |
| CHS.3273 | ENSG00000143630.10 | HCN3 | 41292.75 | 0.59074196 |
| CHS.40544 | ENSG00000196620.10 | UGT2B15 | 40695.96 | 0.60413614 |
| CHS.42530 | ENSG00000133401.16 | PDZD2 | 40638.36 | 0.66326643 |
| CHS.33402 | ENSG00000061656.11 | SPAG4 | 35906.45 | 0.61656889 |
| CHS.29880 | ENSG00000187605.16 | TET3 | 35068.48 | 0.53114392 |
| CHS.2378 | ENSG00000187733.7 | AMY1C | 32857.09 | 1 |
| CHS.18706 | ENSG00000162004.1 | CCDC78 | 32392.95 | 0.6386312 |

| | | | | |
|---|---|---|---|---|
| | 9 | | | 5 |
| CHS.54735 | ENSG00000181085.15 | MAPK15 | 31475.14 | 0.75858312 |
| CHS.35970 | ENSG00000234965.3 | SHISA8 | 31139.08 | 0.85722539 |
| CHS.33642 | ENSG00000124116.19 | WFDC3 | 30772.4 | 0.73865379 |
| CHS.39727 | ENSG00000109758.9 | HGFAC | 30751.99 | 0.59338046 |
| CHS.680 | ENSG00000117245.13 | KIF17 | 30733.07 | 0.54480434 |
| CHS.33331 | ENSG00000131059.12 | BPIFA3 | 30014.13 | 0.53621344 |
| CHS.2070 | ENSG00000055732.13 | MCOLN3 | 29328.39 | 0.50800027 |
| CHS.1453 | ENSG00000198520.12 | ARMH1 | 27883.58 | 0.64666194 |
| CHS.54058 | ENSG00000104450.13 | SPAG1 | 27726.94 | 0.64906405 |
| CHS.17608 | ENSG00000246922.10 | UBAP1L | 27691.07 | 0.65507617 |
| CHS.45982 | ENSG00000204428.12 | LY6G5C | 27461.22 | 0.80460191 |
| CHS.37350 | ENSG00000184345.5 | IQCF2 | 27269.83 | 0.76000804 |
| CHS.25712 | ENSG00000187266.14 | EPOR | 27184.26 | 0.65084906 |
| CHS.7715 | ENSG00000069696.7 | DRD4 | 27181.1 | 0.90773258 |
| CHS.10484 | ENSG00000080854.16 | IGSF9B | 26329.42 | 0.52978417 |
| CHS.39191 | ENSG00000213139.8 | CRYGS | 26202.01 | 0.95179339 |
| CHS.30434 | ENSG00000196862.10 | RGPD4 | 26070.45 | 0.94715396 |
| CHS.39610 | ENSG00000133256.13 | PDE6B | 25470.83 | 0.53342942 |
| CHS.53289 | ENSG00000029534.21 | ANK1 | 25063.51 | 0.62924028 |
| CHS.3444 | ENSG00000243284.1 | VSIG8 | 22802.7 | 0.59956365 |
| CHS.33359 | ENSG00000101440.10 | ASIP | 21793.83 | 0.57223719 |

| | | | | |
|---|---|---|---|---|
| CHS.11737 | ENSG00000135409.11 | AMHR2 | 20896.68 | 0.78936798 |
| CHS.1184 | ENSG00000116819.9 | TFAP2E | 20710.29 | 0.71102626 |
| CHS.18737 | ENSG00000095917.14 | TPSD1 | 20680.31 | 0.91173488 |
| CHS.29099 | ENSG00000189350.13 | TOGARAM2 | 20186.89 | 0.95839082 |
| CHS.23082 | ENSG00000154263.18 | ABCA10 | 19220.93 | 0.9209622 |
| CHS.3248 | ENSG00000163354.15 | DCST2 | 18817.33 | 0.83716872 |
| CHS.13537 | ENSG00000102678.7 | FGF9 | 17834.88 | 0.57370389 |
| CHS.46313 | ENSG00000124602.10 | UNC5CL | 17319.55 | 0.53093065 |
| CHS.30415 | ENSG00000153165.19 | RGPD3 | 16967.64 | 0.84455823 |
| CHS.16888 | ENSG00000186399.11 | GOLGA8R | 16910.95 | 0.76555841 |
| CHS.126 | ENSG00000169885.10 | CALML6 | 16828.01 | 0.68861024 |
| CHS.46363 | ENSG00000112599.9 | GUCA1B | 16793.71 | 0.89168504 |
| CHS.2529 | ENSG00000085465.13 | OVGP1 | 16602.28 | 0.68803381 |
| CHS.33376 | ENSG00000078814.18 | MYH7B | 16276.42 | 0.97451282 |
| CHS.37032 | ENSG00000244607.7 | CCDC13 | 15610.05 | 0.81174692 |
| CHS.53735 | ENSG00000091656.19 | ZFHX4 | 15594.76 | 0.55367059 |
| CHS.10172 | ENSG00000172367.16 | PDZD3 | 14876.57 | 0.61830852 |
| CHS.55898 | ENSG00000197506.8 | SLC28A3 | 14195.73 | 0.71164075 |
| CHS.26738 | ENSG00000160396.9 | HIPK4 | 14172.04 | 0.5215029 |
| CHS.37037 | ENSG00000144648.16 | ACKR2 | 13873 | 0.60777337 |
| CHS.26698 | ENSG00000090932.11 | DLL3 | 13640.57 | 0.51481646 |
| CHS.2091 | ENSG00000171502.15 | COL24A1 | 13221.51 | 0.50062512 |

| | | | | |
|---|---|---|---|---|
| CHS.8830 | ENSG00000149534.9 | MS4A2 | 12956.12 | 0.6051997 |
| CHS.25941 | ENSG00000186526.13 | CYP4F8 | 12573.89 | 0.84151921 |
| CHS.1632 | ENSG00000162383.13 | SLC1A7 | 12557.61 | 0.99671912 |
| CHS.27188 | ENSG00000142233.14 | NTN5 | 12269.62 | 0.79893591 |
| CHS.25809 | ENSG00000161860.8 | SYCE2 | 12233.31 | 0.50840124 |
| CHS.4580 | ENSG00000196660.12 | SLC30A10 | 12123.4 | 0.52622862 |
| CHS.14680 | ENSG00000041515.16 | MYO16 | 11836.23 | 0.93559689 |
| CHS.46033 | ENSG00000241404.7 | EGFL8 | 11823.9 | 0.6341554 |
| CHS.38089 | ENSG00000163424.9 | TEX55 | 11522.05 | 0.51295646 |
| CHS.10181 | ENSG00000235718.9 | MFRP | 11195.52 | 0.9912465 |
| CHS.10173 | ENSG00000248712.8 | CCDC153 | 11114.25 | 0.5362899 |
| CHS.3554 | ENSG00000162746.15 | FCRLB | 11073.19 | 0.91119903 |
| CHS.29911 | ENSG00000144045.14 | DQX1 | 11024.6 | 0.69870744 |
| CHS.36071 | ENSG00000056487.16 | PHF21B | 10952.33 | 0.64446469 |
| CHS.35965 | ENSG00000167077.13 | MEI1 | 10799.88 | 0.53100868 |
| CHS.46405 | ENSG00000146215.14 | CRIP3 | 10699.97 | 0.57025487 |
| CHS.39164 | ENSG00000073803.14 | MAP3K13 | 9752.46 | 0.6076754 |
| CHS.2485 | ENSG00000156150.9 | ALX3 | 9727.31 | 0.78151411 |
| CHS.33264 | ENSG00000180383.4 | DEFB124 | 9525.17 | 0.99896485 |
| CHS.16893 | ENSG00000178115.12 | GOLGA8Q | 9409.95 | 0.736141 |
| CHS.1490 | ENSG00000159588.15 | CCDC17 | 9227.32 | 0.74763312 |
| CHS.6494 | ENSG00000156042.19 | CFAP70 | 9220.08 | 1 |
| CHS.24423 | ENSG00000197705.10 | KLHL14 | 9029.28 | 0.64828314 |
| CHS.23416 | ENSG00000187775.1 | DNAH17 | 8817.94 | 0.6936506 |

| | | | | |
|---|---|---|---|---|
| | 7 | | | 7 |
| CHS.35428 | ENSG00000244752.3 | CRYBB2 | 8719.59 | 0.54323082 |
| CHS.16861 | ENSG00000188626.7 | GOLGA8M | 8278.45 | 0.66455556 |
| CHS.51410 | ENSG00000106384.12 | MOGAT3 | 8258.8 | 0.61698794 |
| CHS.58366 | ENSG00000126733.22 | DACH2 | 8198.81 | 0.87449032 |
| CHS.36738 | ENSG00000183960.9 | KCNH8 | 8120.38 | 0.65598162 |
| CHS.25168 | ENSG00000116032.5 | GRIN3B | 8076.65 | 0.90848557 |
| CHS.42163 | ENSG00000206077.14 | ZDHHC11B | 8011.4 | 0.94585715 |
| CHS.33062 | ENSG00000089101.19 | CFAP61 | 7931.28 | 0.97376212 |
| CHS.26722 | ENSG00000187187.14 | ZNF546 | 7865.49 | 0.55329929 |
| CHS.45731 | ENSG00000112812.16 | PRSS16 | 7678.16 | 0.99967961 |
| CHS.8857 | ENSG00000149506.12 | ZP1 | 7133.54 | 0.68030879 |
| CHS.26643 | ENSG00000182472.9 | CAPN12 | 6788.47 | 0.54544839 |
| CHS.31929 | ENSG00000178568.15 | ERBB4 | 6735.97 | 0.52372709 |
| CHS.17323 | ENSG00000104043.15 | ATP8B4 | 6661.75 | 0.83571584 |
| CHS.31266 | ENSG00000169432.19 | SCN9A | 6597.75 | 0.51330529 |
| CHS.1950 | ENSG00000162620.16 | LRRIQ3 | 6272.54 | 0.58678781 |
| CHS.16855 | ENSG00000153684.16 | GOLGA8F | 6236.75 | 0.96312342 |
| CHS.37435 | ENSG00000157388.20 | CACNA1D | 6070.56 | 0.5092858 |
| CHS.737 | ENSG00000227868.7 | TEX46 | 6024.89 | 0.79268667 |
| CHS.42861 | ENSG00000164512.18 | ANKRD55 | 5922.88 | 0.56186349 |
| CHS.13795 | ENSG00000120669.16 | SOHLH2 | 5887.93 | 0.56105626 |

| | | | | |
|---|---|---|---|---|
| CHS.1690 | ENSG00000162398.12 | LEXM | 5600.67 | 0.60079598 |
| CHS.55350 | ENSG00000186638.17 | KIF24 | 5524.43 | 0.5926711 |
| CHS.6150 | ENSG00000165383.12 | LRRC18 | 4967.56 | 0.82174951 |
| CHS.15935 | ENSG00000119608.15 | PROX2 | 4725.44 | 0.99927203 |
| CHS.36744 | ENSG00000183977.14 | PP2D1 | 4649.83 | 0.58632681 |
| CHS.26732 | ENSG00000105219.10 | CCNP | 4564.82 | 0.74689911 |
| CHS.31617 | ENSG00000151687.15 | ANKAR | 4499.18 | 0.89344725 |
| CHS.8634 | ENSG00000165923.17 | AGBL2 | 4497.62 | 0.89071331 |
| CHS.42139 | ENSG00000153404.15 | PLEKHG4B | 4275.97 | 0.72488815 |
| CHS.46381 | ENSG00000171611.10 | PTCRA | 4243.35 | 0.93241425 |
| CHS.51437 | ENSG00000260097.3 | SPDYE6 | 4179.36 | 0.71972264 |
| CHS.18429 | ENSG00000140470.15 | ADAMTS17 | 4107.88 | 0.9095811 |
| CHS.48122 | ENSG00000125337.21 | KIF25 | 4090.8 | 0.625154 |
| CHS.47740 | ENSG00000118492.18 | ADGB | 3989.99 | 0.77433277 |
| CHS.32359 | ENSG00000237412.7 | PRSS56 | 3915.28 | 0.9972237 |
| CHS.30200 | ENSG00000174501.15 | ANKRD36C | 3760.08 | 0.91833684 |
| CHS.7635 | ENSG00000214279.13 | SCART1 | 3665.92 | 1 |
| CHS.49852 | ENSG00000215045.9 | GRID2IP | 3585.62 | 0.84973589 |
| CHS.10361 | ENSG00000283703.3 | VSIG10L2 | 3369.51 | 1 |
| CHS.34177 | ENSG00000101203.17 | COL20A1 | 3237.8 | 0.51134103 |
| CHS.9594 | ENSG00000149256.16 | TENM4 | 3209.14 | 0.77813682 |
| CHS.30251 | ENSG00000135976.21 | ANKRD36 | 3190.62 | 0.99096414 |
| CHS.39689 | ENSG00000130997.1 | POLN | 2892.54 | 0.5928422 |

| | | | | |
|---|---|---|---|---|
| | 6 | | | 8 |
| CHS.10747 | ENSG00000047617.17 | ANO2 | 2751.73 | 0.97340582 |
| CHS.57205 | ENSG00000204001.10 | LCN8 | 2644.29 | 0.89343831 |
| CHS.10877 | ENSG00000171847.11 | FAM90A1 | 2604.07 | 0.67226687 |
| CHS.1910 | ENSG00000033122.21 | LRRC7 | 2595.13 | 0.52971527 |
| CHS.41521 | ENSG00000250673.3 | REELD1 | 2454.03 | 0.89363211 |
| CHS.52562 | ENSG00000183117.20 | CSMD1 | 2398.11 | 0.75158771 |
| CHS.38404 | ENSG00000170819.5 | BFSP2 | 2345.35 | 0.99898949 |
| CHS.6894 | ENSG00000095587.9 | TLL2 | 2217.06 | 0.8407666 |
| CHS.58272 | ENSG00000225396.6 | FAM236D | 2215.84 | 0.9180356 |
| CHS.38689 | ENSG00000214237.11 | MINDY4B | 2212.09 | 1 |
| CHS.4375 | ENSG00000197721.17 | CR1L | 2178.57 | 0.85584122 |
| CHS.32158 | ENSG00000124003.13 | MOGAT1 | 2029.27 | 0.9988518 |
| CHS.25875 | ENSG00000171136.7 | RLN3 | 2000.65 | 0.91624722 |
| CHS.32093 | ENSG00000243910.8 | TUBA4B | 1996.18 | 0.97624964 |
| CHS.6992 | ENSG00000075891.23 | PAX2 | 1991.34 | 0.53843643 |
| CHS.27340 | ENSG00000142513.6 | ACP4 | 1883.97 | 0.99855624 |
| CHS.35270 | ENSG00000187905.11 | LRRC74B | 1866.63 | 0.99758388 |
| CHS.53259 | ENSG00000188676.15 | IDO2 | 1849.53 | 0.94886809 |
| CHS.6669 | ENSG00000148602.6 | LRIT1 | 1695.28 | 0.91700486 |
| CHS.165344 | ENSG00000284844.2 | TOMT | 1690.05 | 0.80599391 |
| CHS.27346 | ENSG00000174562.14 | KLK15 | 1654.4 | 0.65961074 |
| CHS.43191 | ENSG00000189045.15 | ANKDD1B | 1512.45 | 0.52612648 |

| | | | | |
|---|---|---|---|---|
| CHS.9289 | ENSG00000167800.10 | TBX10 | 1481.57 | 0.95084944 |
| CHS.27218 | ENSG00000268655.3 | ENSG00000268655 | 1453.3 | 0.99094475 |
| CHS.31419 | ENSG00000174279.5 | EVX2 | 1363.45 | 0.56568264 |
| CHS.57259 | ENSG00000198569.10 | SLC34A3 | 1310.65 | 0.55529699 |
| CHS.19997 | ENSG00000070729.14 | CNGB1 | 1284.9 | 0.99701144 |
| CHS.51474 | ENSG00000170615.15 | SLC26A5 | 1275.64 | 0.6725722 |
| CHS.34195 | ENSG00000125508.4 | SRMS | 1273.32 | 0.91842585 |
| CHS.11542 | ENSG00000125084.13 | WNT1 | 1270.01 | 0.6182471 |
| CHS.54737 | ENSG00000203499.12 | IQANK1 | 1263.05 | 1 |
| CHS.51163 | ENSG00000240720.10 | LRRD1 | 1182.12 | 0.85101343 |
| CHS.41989 | ENSG00000187821.9 | HELT | 1135.26 | 0.76704896 |
| CHS.46904 | ENSG00000164411.12 | GJB7 | 1089.98 | 0.98005468 |
| CHS.16907 | ENSG00000134160.15 | TRPM1 | 1076.05 | 0.59275127 |
| CHS.53249 | ENSG00000169495.5 | HTRA4 | 1074.16 | 0.5211421 |
| CHS.46515 | ENSG00000124818.15 | OPN5 | 1047 | 0.91209169 |
| CHS.25553 | ENSG00000142449.13 | FBN3 | 1003.27 | 0.52324898 |

**Table S2. Gene counts of each biotype in CHESS 3 annotation on GRCh38 vs. CHM13.**

| **Gene biotype** | **Number of genes in CHESS3** | **Number mapped onto CHM13** |
|---|---|---|
| protein coding | 19839 | 19968 |
| lncRNA | 17623 | 18611 |
| pseudogene | 16572 | 17405 |
| miRNA | 1914 | 2208 |
| snoRNA | 1195 | 1188 |

| | | |
|---|---|---|
| rRNA | 40 | 765 |
| tRNA | 453 | 552 |
| VDJ segments | 565 | 543 |
| snRNA | 153 | 192 |
| misc RNA | 35 | 182 |
| ncRNA | 28 | 28 |
| TEC | 27 | 30 |
| C region/C region pseudogene | 26 | 29 |
| antisense RNA | 19 | 19 |
| other | 12 | 12 |
| vault RNA | 4 | 4 |
| scRNA | 4 | 4 |
| Y RNA | 4 | 7 |
| telomerase RNA | 1 | 1 |
| ncRNA pseudogene | 1 | 1 |
| RNase P RNA | 1 | 1 |
| RNase MRP RNA | 1 | 1 |

**Table S3. Variant counts in CHESS 3 annotation of CHM13 vs. GRCh38 predicted by LiftoffTools.**

| **Variant effect** | **Number of events** | **Number of genes** |
|---|---|---|
| nonsynonymous | 18800 | 4916 |
| synonymous | 15370 | 3444 |
| inframe deletion | 534 | 164 |
| inframe insertion | 506 | 169 |
| frameshift | 454 | 236 |
| stop gained | 176 | 91 |
| start lost | 127 | 59 |
| 3’ truncated | 10 | 8 |
| 5’ truncated | 1 | 1 |

**Supplementary Table S4.** Discarded GTEx samples. 133 samples were outliers due to significantly lower than average alignment rate, fewer reads, and/or excessively variable read length. Samples listed here were not used in any part of the analysis. The number of reads in the discarded samples varied from 2,548 to 1,190,719 ,while the 1st and 3rd quartiles across all GTEx samples used in this study were 37,224,913 and 51,526,165 respectively. Similarly, the alignment rate observed in the discarded samples was between 3.57% and 72.58% compared to

| the 89.00% and 93.25% for the 1st and 3rd quartiles respectively for the full data set. | | | |
|---|---|---|---|
| **Tissue** | **Sample** | **# of Reads** | **% Alignment** |
| Adipose Tissue | SRR2135413 | 467460 | 65.34 |
| Adrenal Gland | SRR2135287 | 889264 | 49.04 |
| Adrenal Gland | SRR2135339 | 132582 | 36.52 |
| Bladder | SRR2135324 | 550518 | 51.53 |
| Bladder | SRR2135407 | 763831 | 48.67 |
| Blood Vessel | SRR2135290 | 753835 | 37.85 |
| Blood Vessel | SRR2135291 | 913004 | 34.7 |
| Blood Vessel | SRR2135328 | 860470 | 45.13 |
| Blood Vessel | SRR2135348 | 458578 | 59.51 |
| Brain | SRR2135333 | 861593 | 68.27 |
| Brain | SRR2135334 | 618537 | 67.91 |
| Brain | SRR2135335 | 767618 | 53.16 |
| Brain | SRR2135336 | 522191 | 60.07 |
| Brain | SRR2135337 | 585061 | 64.62 |
| Brain | SRR2135338 | 803585 | 63.43 |
| Brain | SRR2135341 | 721241 | 54.05 |
| Brain | SRR2135342 | 894945 | 65.14 |
| Brain | SRR2135343 | 787035 | 62.64 |
| Brain | SRR2135355 | 709235 | 69.18 |
| Brain | SRR2135356 | 564478 | 70.4 |
| Brain | SRR2135358 | 655355 | 60.36 |
| Brain | SRR2135359 | 344967 | 63.88 |
| Brain | SRR2135360 | 628712 | 60.7 |
| Brain | SRR2135361 | 747160 | 49.26 |
| Brain | SRR2135363 | 797891 | 67.76 |
| Brain | SRR2135374 | 752192 | 72.58 |
| Breast | SRR2135303 | 545813 | 62.64 |
| Breast | SRR2135304 | 829860 | 38.66 |
| Breast | SRR2135332 | 51521 | 25.15 |
| Breast | SRR2135370 | 554418 | 57.01 |
| Colon | SRR2135297 | 773172 | 69.2 |
| Colon | SRR2135298 | 741198 | 41.66 |
| Colon | SRR2135299 | 820420 | 56.06 |

| | | | |
|---|---|---|---|
| Colon | SRR2135315 | 1014825 | 39.39 |
| Colon | SRR2135325 | 641420 | 45.03 |
| Colon | SRR2135395 | 850148 | 48.99 |
| Colon | SRR2135397 | 798490 | 48.36 |
| Colon | SRR2135398 | 602904 | 49.97 |
| Colon | SRR2135414 | 753297 | 45.54 |
| Colon | SRR2135416 | 345269 | 66.01 |
| Esophagus | SRR2135293 | 762958 | 48.12 |
| Esophagus | SRR2135294 | 945378 | 44.67 |
| Esophagus | SRR2135313 | 730734 | 56.95 |
| Esophagus | SRR2135327 | 81723 | 38.25 |
| Esophagus | SRR2135329 | 565980 | 45.54 |
| Esophagus | SRR2135350 | 600065 | 55.82 |
| Esophagus | SRR2135352 | 39361 | 29.28 |
| Esophagus | SRR2135362 | 729270 | 40.19 |
| Esophagus | SRR2135367 | 738297 | 48.42 |
| Esophagus | SRR2135368 | 813384 | 60.58 |
| Esophagus | SRR2135369 | 631816 | 41.68 |
| Esophagus | SRR2135381 | 616924 | 44.78 |
| Esophagus | SRR2135382 | 837370 | 37.96 |
| Esophagus | SRR2135393 | 1131576 | 34.87 |
| Esophagus | SRR2135394 | 899464 | 46.71 |
| Esophagus | SRR2135409 | 818462 | 50.79 |
| Esophagus | SRR2135410 | 621391 | 46.29 |
| Esophagus | SRR2135411 | 666012 | 54.53 |
| Heart | SRR2135289 | 858262 | 29.21 |
| Heart | SRR2135310 | 926101 | 32.38 |
| Heart | SRR2135347 | 761241 | 51.03 |
| Heart | SRR2135392 | 1025710 | 46.19 |
| Kidney | SRR2135353 | 769043 | 53.25 |
| Kidney | SRR2135396 | 608904 | 61.17 |
| Liver | SRR2135317 | 775788 | 47.8 |
| Liver | SRR2135349 | 909967 | 32.68 |
| Liver | SRR2135365 | 439200 | 60.53 |
| Liver | SRR2135366 | 606470 | 17.78 |
| Liver | SRR2135380 | 756247 | 39.42 |

| | | | |
|---|---|---|---|
| Liver | SRR2135387 | 820528 | 38.77 |
| Lung | SRR2135292 | 489724 | 57.77 |
| Lung | SRR2135311 | 452184 | 69.38 |
| Lung | SRR2135346 | 563037 | 61.97 |
| Lung | SRR2135351 | 789936 | 68.19 |
| Lung | SRR2135364 | 496368 | 62.04 |
| Lung | SRR2135379 | 539538 | 67.39 |
| Lung | SRR2135391 | 420732 | 67.53 |
| Muscle | SRR2135308 | 424465 | 67.27 |
| Muscle | SRR2135331 | 505377 | 63.54 |
| Muscle | SRR2135378 | 610879 | 62.05 |
| Ovary | SRR2135322 | 59731 | 30.15 |
| Ovary | SRR2135371 | 610971 | 49.1 |
| Ovary | SRR2135400 | 705859 | 43.1 |
| Ovary | SRR2135405 | 620557 | 59.57 |
| Pancreas | SRR2135288 | 643775 | 51.2 |
| Pancreas | SRR2135296 | 817930 | 41.5 |
| Pancreas | SRR2135312 | 727517 | 43.12 |
| Pancreas | SRR2135318 | 569411 | 65.65 |
| Pancreas | SRR2135383 | 862339 | 63.73 |
| Pancreas | SRR2135388 | 916474 | 48.89 |
| Pituitary | SRR2135357 | 2548 | 49.2 |
| Pituitary | SRR2135375 | 750912 | 51.48 |
| Pituitary | SRR2135376 | 703044 | 50.76 |
| Prostate | SRR2135300 | 663930 | 47.4 |
| Prostate | SRR2135316 | 782713 | 41.65 |
| Prostate | SRR2135386 | 502325 | 65.06 |
| Salivary Gland | SRR2135402 | 664055 | 45.68 |
| Skin | SRR2135286 | 940964 | 27.14 |
| Skin | SRR2135305 | 569414 | 66.24 |
| Skin | SRR2135306 | 550000 | 60.76 |
| Skin | SRR2135319 | 717304 | 66.56 |
| Skin | SRR2135321 | 431526 | 62.39 |
| Skin | SRR2135417 | 571327 | 57.69 |
| Small Intestine | SRR2135399 | 786667 | 46.79 |
| Small_Intestine | SRR2135415 | 155918 | 68.04 |

| | | | |
|---|---|---|---|
| Spleen | SRR2135307 | 448958 | 38.55 |
| Spleen | SRR2135309 | 904438 | 48.45 |
| Spleen | SRR2135389 | 751864 | 49.35 |
| Stomach | SRR2135295 | 734465 | 41.04 |
| Stomach | SRR2135314 | 1190719 | 3.57 |
| Stomach | SRR2135323 | 824325 | 34.06 |
| Stomach | SRR2135373 | 773853 | 51.87 |
| Stomach | SRR2135384 | 854838 | 31.07 |
| Stomach | SRR2135385 | 779252 | 59.66 |
| Stomach | SRR2135412 | 568688 | 60.38 |
| Testis | SRR2135285 | 625506 | 50.12 |
| Testis | SRR2135301 | 542106 | 64.09 |
| Testis | SRR2135302 | 678632 | 58.79 |
| Testis | SRR2135320 | 509297 | 70.02 |
| Testis | SRR2135354 | 44068 | 35.68 |
| Testis | SRR2135377 | 550160 | 56.34 |
| Thyroid | SRR2135283 | 1097525 | 34.41 |
| Thyroid | SRR2135284 | 1004719 | 17.98 |
| Thyroid | SRR2135326 | 662069 | 64.6 |
| Thyroid | SRR2135344 | 582791 | 65.43 |
| Thyroid | SRR2135345 | 638187 | 46.58 |
| Thyroid | SRR2135390 | 629065 | 65.44 |
| Thyroid | SRR2135403 | 557144 | 64.52 |
| Uterus | SRR2135330 | 648893 | 42.45 |
| Uterus | SRR2135372 | 851435 | 38.57 |
| Uterus | SRR2135406 | 793552 | 52.19 |
| Vagina | SRR2135401 | 1059781 | 44.79 |
| Vagina | SRR2135408 | 816997 | 46.18 |
